# Loss of mitochondrial protease CLPP activates type I interferon responses through the mtDNA-cGAS-STING signaling axis

**DOI:** 10.1101/2020.08.30.274712

**Authors:** Sylvia Torres-Odio, Yuanjiu Lei, Suzana Gispert, Antonia Maletzko, Jana Key, Saeed Menissy, Ilka Wittig, Georg Auburger, A. Phillip West

## Abstract

Caseinolytic mitochondrial matrix peptidase proteolytic subunit, CLPP, is a serine protease that degrades damaged or misfolded mitochondrial proteins. CLPP null mice exhibit growth retardation, deafness, and sterility, resembling human Perrault syndrome (PS), but also display immune system alterations. However, the molecular mechanisms and signaling pathways underlying immunological changes in CLPP null mice remain unclear. Here we report the steady state activation of type I interferon (IFN-I) signaling and antiviral gene expression in CLPP deficient cells and tissues. Depletion of the cyclic GMP-AMP (cGAS)-Stimulator of Interferon Genes (STING) DNA sensing pathway ablates heightened IFN-I responses and abrogates the broad viral resistance phenotype of CLPP null cells. Moreover, we report that CLPP deficiency leads to mitochondrial DNA (mtDNA) instability and packaging alterations. Pharmacological and genetic approaches to deplete mtDNA or inhibit cytosolic release markedly reduce antiviral gene expression, implicating mtDNA stress as the driver of IFN-I signaling in CLPP null mice. Our work places the cGAS-STING-IFN-I innate immune pathway downstream of CLPP and may have implications for understanding myriad human diseases involving CLPP dysregulation.

## Introduction

Mitochondrial proteases are key modulators of several mitochondrial functions, including the maturation of proteins, maintenance of protein quality control, control of mitochondrial gene expression and biogenesis, mitophagy, and apoptosis (Quirós et al., 2015). CLPP, caseinolytic mitochondrial matrix peptidase proteolytic subunit, is a highly conserved, processive serine protease located in the mitochondrial matrix. CLPP is the protease component of the CLPXP complex that cleaves peptides and various proteins in an ATP-dependent process, together with chaperone and ATPase CLPX (Baker and Sauer, 2012). While prokaryotic ClpXP has been extensively studied, the role and function of CLPXP in mitochondria remain less clear, and there is limited information regarding its specific substrates (Bhandari et al., 2018). Studies in *C. elegans* have implicated CLPXP as a key component of the mitochondrial unfolded protein response (UPR^mt^), and downstream signaling and transcriptional responses of the UPR^mt^ are attenuated in worms lacking CLPP activity (Haynes et al., 2007; Pellegrino et al., 2014). Mammalian CLPP is involved in pleiotropic cellular functions such as myoblast differentiation, cell proliferation, and mitoribosome assembly, which controls the rate of mitochondrial protein synthesis (Deepa et al., 2016; Szczepanowska et al., 2016). Moreover, CLPP has recently been implicated in cancer, and hyperactivating CLPP was shown to alter mitochondrial function and selectively kill some cancer cells (Ishizawa et al., 2019). Thus, CLPP is being studied as a biological target of anti-cancer drugs in current clinical trials (Graves et al., 2019).

Studies performed in CLPP null mice (CLPP-KO) highlight the physiological and pathological relevance of this model to the human disorder known as Perrault syndrome (PS). CLPP-KO mice display profound phenotypic changes characterized by growth retardation, deafness, and premature sterility, despite exhibiting only mild mitochondrial bioenergetic deficits (Gispert et al., 2013). Similar phenotypes are also observed in PS, where the genetic component involves autosomal recessive mutations in CLPP and other nuclear DNA-encoded mitochondrial proteins (Jenkinson et al., 2013). Other phenotypes of this mouse include insulin resistance and protection from diet-induced obesity (Bhaskaran et al., 2018), accelerated depletion of ovarian follicular reserve (T. Wang et al., 2018), and impairment of adaptive thermogenesis due to a decline in brown adipocytes (Becker et al., 2018).

Interestingly, CLPP-KO mice show activation of antiviral genes, with elevated expression of interferon stimulated genes (ISGs) noted in many tissues and fibroblasts (Gispert et al., 2013; Key et al., 2020). Activation of innate immune signaling downstream of mitochondrial dysfunction has been reported in several mitochondrial mutant animals (Pellegrino et al., 2014; West et al., 2015), as well as in animal models of human disease and patient samples (Nakahira et al., 2015; West, 2017; West and Shadel, 2017). Emerging evidence localizes cGAS, Toll-like receptor 9 (TLR9), and the NOD-, LRR- and pyrin domain-containing protein 3 (NLRP3) inflammasome downstream of mitochondrial damage (Nakahira et al., 2015; West, 2017; West and Shadel, 2017).

Eukaryotic mitochondria maintain prokaryotic features, including a double-stranded circular DNA genome, inner membrane cardiolipin, and N-formylated proteins, which act as potent triggers of cGAS, TLR9, NLRP3, and other innate immune sensors when released from mitochondria into the cytosol or extracellular space. The release and accumulation of these so-called mitochondrial called damage-associated molecular patterns (DAMPs) is increasingly implicated in the inflammatory pathology of numerous human disorders including autoimmunity, neurodegenerative diseases, and cancer (Nakahira et al., 2015; West and Shadel, 2017; Youle, 2019). The mitochondrial molecular mechanisms and innate immune signaling pathways underlying activation of antiviral signatures in CLPP-KO mice remain uncharacterized, and the biological significance of this phenotype is unclear. Moreover, the role of IFN-I signaling in CLPP-KO mouse pathology and its possible implication in human PS remain to be examined.

Herein we show that enhanced ISG expression and IFN-I responses in CLPP-KO mice are triggered by mtDNA stress and mediated by the cGAS-STING DNA sensing pathway. Deletion of STING or the interferon α/β receptor (IFNAR) in CLPP-KO MEFs ablates steady state expression of broadly antiviral ISGs and reduces the potent antiviral phenotypes observed during RNA and DNA virus infection. These findings shed light on a new mitochondrial pathway upstream of cGAS and STING, link mitochondrial proteostasis to mtDNA genome maintenance, and may have important implications for understanding mitochondrial-innate immune crosstalk in the context of human health and disease.

## Results and Discussion

### CLPP deficiency leads to steady state ISG expression and an antiviral signature in cells and tissues

We first expanded upon our prior findings that revealed increased steady state expression of antiviral genes in CLPP deficient tissues (Gispert et al., 2013). Using Ingenuity Pathway Analysis (IPA) software, we observed that predicted upstream regulators of the antiviral gene signature in CLPP-KO tissues included factors governing IFN-I signaling (*Irf7*, *Irf3* and *Stat1)*, IFNα, and IFNAR (Fig EV1A). Among the list of upregulated IRF7 and STAT1 target genes, we identified canonical ISGs such as *Usp18*, *Ifit1b*, *Ifi44*, and *Rtp4* (Fig EV1B). Individual analyses of heart and liver tissues from 3-5-month-old mice revealed a significant enhancement of ISGs at the protein level in both the liver and heart of CLPP-KO mice (Fig EV1C and D).

To better characterize this ISG response, we used mouse embryonic fibroblasts (MEFs) and human foreskin fibroblasts deficient in CLPP. Expression profiles of CLPP-KO MEFs revealed an enrichment of ISGs and antiviral signaling factors at both the RNA and protein level (Fig 1A and B). Among the set of overexpressed ISGs, we observed those involved in direct antiviral activity (VIPERIN/Rsad2, IFIT1, IFIT3); regulators of antiviral and IFN-I signaling (USP18 and STAT1); as well as RNA and DNA sensors (RIG-I and ZBP1). Global proteome profiling revealed that greater than 60 percent of the most differentially expressed proteins in CLPP-KO MEFs were ISGs (Fig 1C). Transfection of CLPP-KO MEFs with poly(I:C), a dsRNA analog agonist of the cytosolic RNA helicase MDA5, or challenge with LPS, a TLR4 ligand, revealed elevated ISG expression compared to WT, which indicates potentiation/priming of antiviral signaling pathways downstream of both cytosolic and membrane-bound pattern recognition receptors (Fig 1D and E). To assess if this heightened ISG response was only observed in mouse cells, we transduced human foreskin fibroblasts with short hairpin RNAs targeting CLPP (shCLPP) or EGFP as a control (shEGFP). Similar to our findings in MEFs and murine tissues, reduction of CLPP in human fibroblasts led to the steady state upregulation of antiviral ISGs at the protein level, including IFIT2, IFITM1, RIG-I, STAT1 and others (Fig 1F).

**Figure 1.**
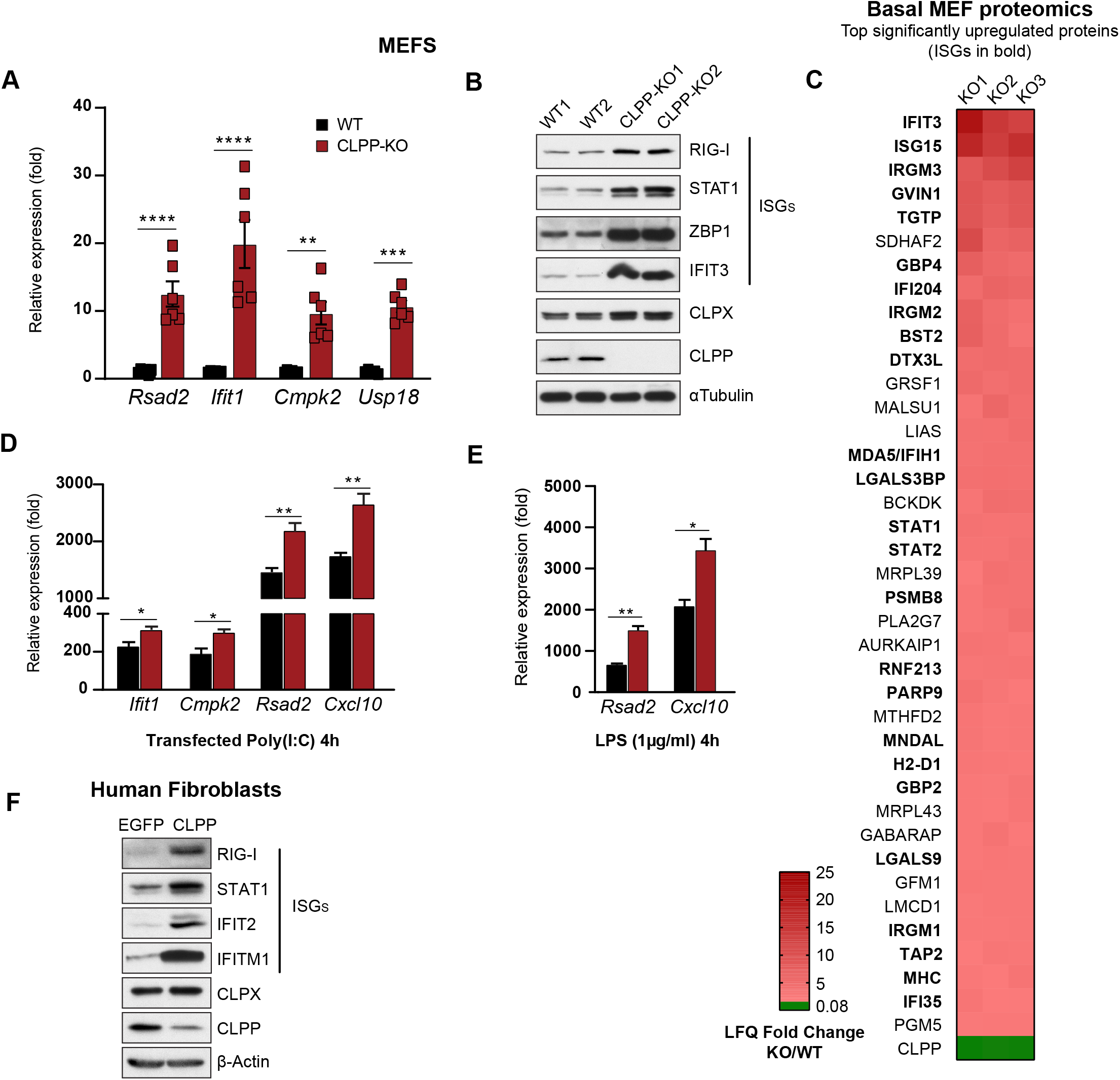
CLPP deficiency leads to steady state interferon stimulated gene (ISG) expression in mouse and human cells. A, B Quantitative real-time PCR (A) and western blots (B) of baseline ISGs of littermate WT and CLPP-KO mouse embryonic fibroblasts lines (n=2). C Heat map from the top 40 significantly (p<0.05) upregulated proteins from baseline proteomic data of littermates WT and CLPP-KO MEFs lines. Data represented as Label-free quantification (LFQ) Fold Change of each CLPP-KO MEFs over average WT (n=3). D, E Quantitative real-time PCR of ISGs in WT and CLPP-KO MEFs, 4h after transfection with poly(I:C) (D) or LPS challenge (E). F Human foreskin fibroblasts were transduced with the shRNA against CLPP and EGFP (as control) and selected with puromycin (2μg/ml). After selection, cells were plated in 12-well dishes and harvested for western blots of baseline ISGs. Data information: Data are presented as mean ± s.e.m. of triplicate technical replicates and are representative of three independent experiments. *p<0.05, **p<0.01, ***p<0.001, ****p<0.0001. In A, two-way ANOVA Tukey–8217;s *post-hoc*. In D, E Student–8217;s t-test

### STING and IFNAR mediate the steady state ISG and antiviral signature observed in CLPP deficient cells and tissues

Recent work has revealed that the release of mitochondrial nucleic acids into the cytosol is a potent trigger of DNA and RNA sensors of the innate immune system (Dhir et al., 2018; Rongvaux et al., 2014; West et al., 2015). To next pinpoint downstream sensors mediating the antiviral response in CLPP-KO cells, we transiently knocked down the DNA sensor cGAS or the cytosolic RNA adaptor MAVS and quantified ISG responses in MEFs. Downregulation of cGAS profoundly diminished the ISG response in CLPP-KO MEFs (Fig 2A) while MAVS depletion did not (Fig. EV2A), implicating the cytosolic DNA sensing machinery as the main trigger of ISGs in CLPP deficient cells. Using a parallel approach to bypass any off-target and immune stimulatory effects of transient siRNA transfection, we crossed CLPP heterozygous mutant mice with *Sting*^*gt//gt*^ and *Ifnar*^*-/-*^ strains to generate CLPP-KO/STING^gt/gt^ and CLPP-KO/IFNAR-KO mice, respectively, and assessed the antiviral response in MEFs and tissues. Consistent with our siRNA data implicating the cGAS pathway, the relative expression of ISGs *Usp18*, *Isg15, and Cxcl10* was markedly reduced in the absence of STING (Fig 2B), with this effect being more pronounced in CLPP-KO/IFNAR-KO MEFs. Ablation of elevated IFN-I signaling was also observed in RNA extracts from tissues of the double mutant crosses, as qRT-PCR analysis of heart (Fig 2C) and liver (Fig EV2B) from double KO mice showed diminished ISG RNA compared to CLPP-KO tissues. A similar effect was seen for antiviral proteins such as RIG-I, STAT1, ZBP1, and IFIT3 in heart extracts (Fig 2D), with marked decreases in all ISGs in CLPP-KO/STING^gt/gt^ and CLPP-KO/IFNAR-KO mice compared to CLPP-KO alone. Collectively, our findings indicate that loss of CLPP in murine cells and tissues leads to exacerbated IFN-I signaling via engagement of the cGAS-STING axis.

**Figure 2.**
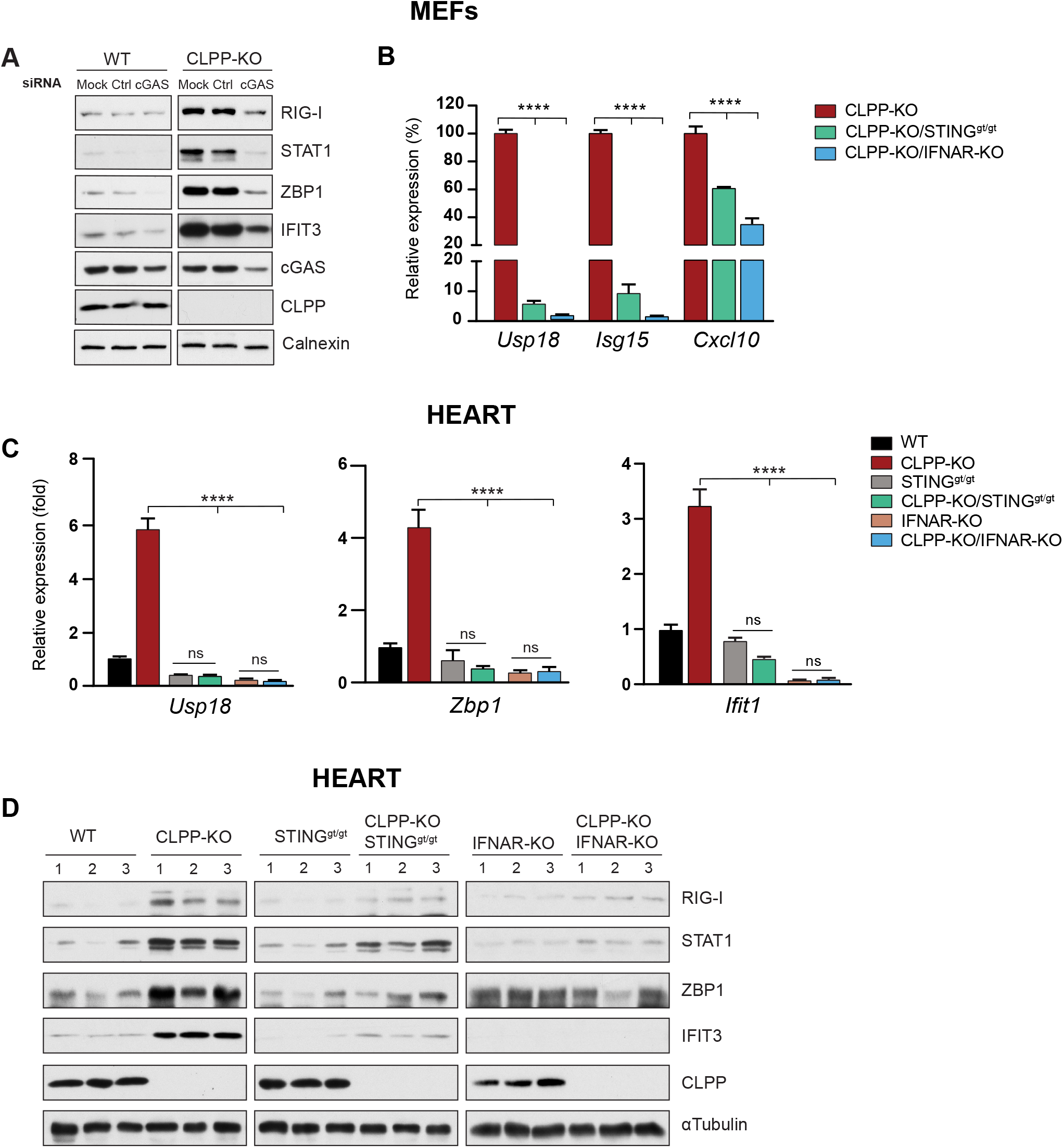
STING and IFNAR mediate the steady state ISG and antiviral signature in CLPP deficient cells and tissues. A Western blots of ISGs in WT and CLPP-KO MEFs transfected with siRNA Control (Ctrl) and sicGAS for 72hrs. B Quantitative real-time PCR of baseline ISGs in WT, CLPP-KO, and CLPP-KO/STING^gt/gt^ or CLPP-KO/IFNAR-KO double mutant MEFs (n=2 MEF lines). Real-time PCR data was represented as relative expression percentage (%) with CLPP-KO set to 100%. C, D Quantitative real-time PCR (C) and western blots (D) of baseline ISGs from heart tissue from 12-month old male mice (n=3). Data information: In B and C data are presented as mean ± s.e.m. of triplicate technical replicates. ***p<0.001, ****p<0.0001, ns, non-significant. (One-way ANOVA, Tukey–8217;s *post-hoc*)

CLPP-KO mice exhibit reduced adiposity, increased whole-body energy expenditure, and protection from diet-induced obesity, indicating a role of CLPP in metabolic rewiring (Bhaskaran et al., 2018). Although ablation of STING and IFNAR was sufficient to blunt the steady state ISGs signature in CLPP-KO cells and tissues, loss of STING or IFNAR signaling was not sufficient to rescue the growth deficits of CLPP-KO mice (Fig EV2C). Moreover, STING depletion did not significantly impact body composition changes (i.e. fat/lean ratio via EchoMRI) resulting from loss of CLPP (Fig EV2D). Our study did not investigate the role of STING-IFN-I signaling in infertility or deafness in CLPP-KO mice, which are also key features of human PS. However, several studies have linked IFN-I to sterility in both transgenic mouse models and human patients receiving exogenous IFN-α therapy (Hekman et al., 1988; Iwakura et al., 1988; Ulusoy et al., 2004). Moreover, sudden hearing loss has been reported as an adverse effect of IFN-α immunotherapy in patients with hepatitis C virus or cancer (Asal et al., 2014.; Formann et al., 2004). Therefore, it will be interesting to explore whether ablation of STING or IFNAR in CLPP-KO mice impacts the development or progression of sterility and/or deafness, which may have implications for understanding the role of STING-IFN-I signaling in the diverse pathology of human PS.

### CLPP-KO cells are more resistant to viral infection owing to elevated activation of the STING pathway

To next assess if basal cGAS-STING-IFN-I signaling and increased steady state expression ISGs in CLPP-KO cells is functionally relevant, we infected MEFs with recombinant Vesicular Stomatitis Virus (VSV) expressing the viral protein VSV-G fused to GFP (VSV-G/GFP) (Dalton and Rose, 2001). After infection at a MOI of 0.1 for 24h, microscopic analysis showed a striking absence of GFP fluorescence in CLPP-KO cells compared to WT controls (Fig 3A). Quantification of GFP (Fig EV3A) indicated that only 1.3% of CLPP-KO cells were positive for GFP compared to 25% of WT cells. STING^gt/gt^ cells were more susceptible to VSV infection consistent with prior results (Ahn and Barber, 2019; Franz et al., 2018), with more than 60% of cells staining GFP positive. Similarly, 68% of CLPP-KO/STING^gt/gt^ cells were GFP positive, indicating that viral resistance of CLPP-KO cells was completely lost in CLPP-KO/STING^gt/gt^ MEFs. Relative levels of viral RNA transcripts for *VSV-G* and *VSV-M* were reduced greater than 95% in CLPP-KO cells compared to WTs (Fig 3B). In contrast, viral transcript levels were equally high in VSV-infected STING^gt/gt^ and CLPP-KO/STING^gt/gt^ cells. Similar results were found at earlier VSV infection timepoints (i.e. 16 hpi, Fig EV3B), corroborating microscopic results demonstrating that antiviral phenotypes of CLPP-KO cells are mediated by STING signaling.

**Figure 3.**
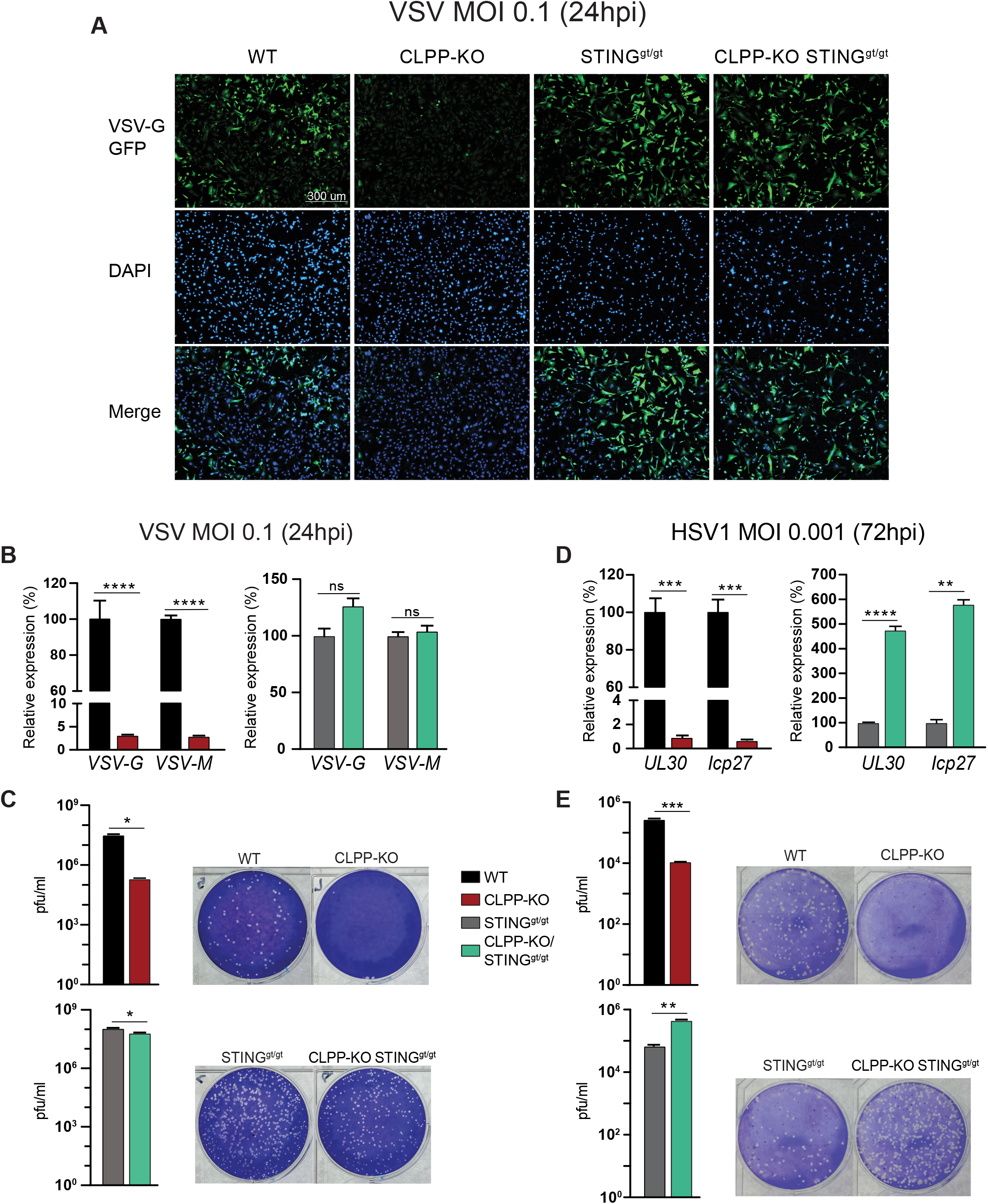
CLPP-KO fibroblasts are more resistant to viral infection owing to elevated activation of the STING pathway. A-C Analysis of viral load and viral plaques in MEFs after 24hrs of VSV infection at multiplicity of infection (MOI) of 0.1. (A) Microscopy images of viral GFP expression in MEFs infected with VSV. DAPI was used for nuclear staining. (B) Quantitative real-time PCR of viral RNA transcripts, *VSV-G* and *VSV-M*, after VSV infection (n=3 MEF lines). (C) Quantification of plaque assays from VSV infected MEFs. D Quantitative real-time PCR of viral RNA in MEFs infected with HSV-1 at 72hrs post infection (hpi) and MOI of 0.001 (n=3 MEF lines). E Quantification of plaque assay from HSV-1 infected MEFs. Plaque figure is representative of 10^−2^ supernatant dilution and graphed as pfu/ml (n=2). Data information: In (C, E), plaque figure is representative of 10^−5^ supernatant dilution and graphed as pfu/ml. In (B, D), Real-time PCR data was represented as relative expression percentage (%) comparing CLPP-KO vs WT and CLPP-KO/STING^gt/gt^ vs STING^gt/gt^ MEFs. In (B-E), data are presented as mean ± s.e.m. of triplicate technical replicates. **p<0.01, ***p<0.001, ****p<0.0001, ns non-significant (Student–8217;s t-test)

To further evaluate the resistance of CLPP-KO cells to viral infection/replication, we employed plaque assays to quantify the level of functional VSV virions released into the supernatant of infected cells (Fig 3C). At 24hpi, there was a striking reduction of viral plaques in CLPP-KO MEFs (2-log reduction) compared to WTs. This contrasted strongly with CLPP-KO/STING^gt/gt^ cells, which exhibited only a small, 1.7-fold reduction compared to STING^gt/gt^ littermate MEFs. The viral protection observed in CLPP-KO cells is likely due to both elevated baseline and induced ISGs, as levels of antiviral genes *Usp18* and *Cmpk2* were further enhanced in CLPP-KO over WT controls after VSV infection (Fig EV3C). Although the VSV-mediated upregulation of ISGs in CLPP-KO cells was greatly blunted by loss of STING, the fold induction of ISGs in double mutant CLPP-KO/STING^gt/gt^ MEFs over infected STING^gt/gt^ MEFs was similar to CLPP-KO over WT cells. This is an expected result considering that RNA viruses such as VSV predominantly engage the RIG-I-MAVS pathway, which remains intact and active in the absence of STING. Thus, the baseline antiviral priming in CLPP-KO cells driven by persistent STING-IFN-I signaling, and not the potentiated induction of ISGs during infection, is likely the most important factor governing the resistance of CLPP-KO cells to VSV.

To determine if the viral resistance extended to DNA viruses, we infected MEF lines with HSV-1, a dsDNA virus that employs several diverse mechanisms to evade innate antiviral responses. After infection at a MOI 0.001 and incubation for 72 hours, HSV-1 viral RNAs *UL30* and *Icp27* were reduced greater than 95% in CLPP-KO MEFs compared to WT controls (Fig 3D), mirroring findings seen with VSV infections. The dramatic reduction in HSV-1 RNA expression in CLPP-KO cells was lost in double KO cells, and in fact we observed that CLPP-KO/STING^gt/gt^ MEFs exhibited five times more viral RNA transcripts than STING^gt/gt^ littermates (Fig 3D). This marked increase in viral RNAs in the double KO cells was not observed in VSV infections, where there was only a minor increase in viral transcripts in CLPP-KO/STING^gt/gt^ MEFs over infected STING^gt/gt^ MEFs (Fig 3C). A similar trend was observed in quantification of viral particles by plaque assays, as we observed a marked resistance of CLPP-KO cells to HSV-1 (1.5 log reduction in plaques) compared to WT cells (Fig 3E). However, CLPP-KO/STING^gt/gt^ cells were more susceptible to HSV-1 infection (0.5 log increase in plaques) compared to STING^gt/gt^, consistent with viral RNA expression data.

We next assessed whether the strong antiviral response was maintained at a higher and more lytic MOI, where HSV-1 virulence and IFN-I-repressive mechanisms are more potent. At a MOI of 0.01 48hpi, CLPP-KO cells remained more resistant to HSV-1, albeit to a lesser extent compared to infection with a lower MOI (Fig EV3D). CLPP-KO cells showed a 40% reduction in *UL30* and *Icp27* expression compared to WTs. However, similar to results from MOI 0.001 infections, CLPP-KO/STING^gt/gt^ MEFs exhibited 5 times more viral RNA than STING^gt/gt^ cells (Fig EV3D). The interesting finding that CLPP-KO/STING^gt/gt^ MEFs are more susceptible to HSV-1 infection may be due to the fact that HSV-1 is a robust modulator of host mitochondrial function and metabolism (Duarte et al., 2019; Reshi et al., 2018). We hypothesize that the complete lack of cytosolic DNA sensing and antiviral IFN-I, combined with altered mitochondrial homeostasis in CLPP-KO/STING^gt/gt^ cells, likely synergize to increase HSV-1 virulence mechanisms and increase viral replication. Notably, HSV-1 can directly target host mtDNA through the virally-encoded deoxyribonuclease UL12.5, resulting in nucleoid stress/enlargement followed by rapid mtDNA depletion (Corcoran et al., 2009; Saffran et al., 2007; West et al., 2015). Thus, increased interference with mtDNA homeostasis by UL12.5 or other encoded factors may potentially explain the elevated susceptibility of double KO cells to HSV-1. In sum, our results show that potentiated cGAS-STING-dependent ISG and IFN-I responses in CLPP-KO cells leads to a robust antiviral state that is broadly restrictive to both RNA and DNA viruses.

### CLPP deficiency results in altered mtDNA abundance and nucleoid morphology

Our prior work has documented that mtDNA stress and cytosolic release is a potent inducer of IFN-I responses via the cGAS-STING DNA sensing axis (West et al., 2015). Since the ISG and antiviral priming phenotypes in CLPP-KO cells and tissues were dependent on cGAS-STING, we hypothesized that mtDNA might be the mitochondrial DAMP triggering this response. Interestingly, we found that MEFs and tissues from CLPP-KO mice exhibit baseline elevations in mtDNA abundance by qPCR-based approaches (Fig EV4A), despite no coordinate upregulation in the mtDNA packaging protein Transcription factor A mitochondrial (TFAM) (Gispert et al., 2013) or full-length mtDNA genomes (Szczepanowska et al., 2016). We reasoned that this might indicate the presence of mtDNA instability in CLPP deficient cells. In agreement with this hypothesis, we observed that CLPP-KO MEFs (Fig 4A) and CLPP-depleted human fibroblasts (Fig EV4B) displayed significant mtDNA nucleoid enlargement and aggregation by confocal microscopy. These results are similar to the mtDNA stress phenotype observed in *Tfam*^*+/−*^ MEFs, which have enlarged mtDNA nucleoids and mitochondrial network hyperfusion that result in the release and cytosolic accumulation of mtDNA, tonic cGAS-STING-IFN-I signaling, and antiviral priming (West et al., 2015).

**Figure 4.**
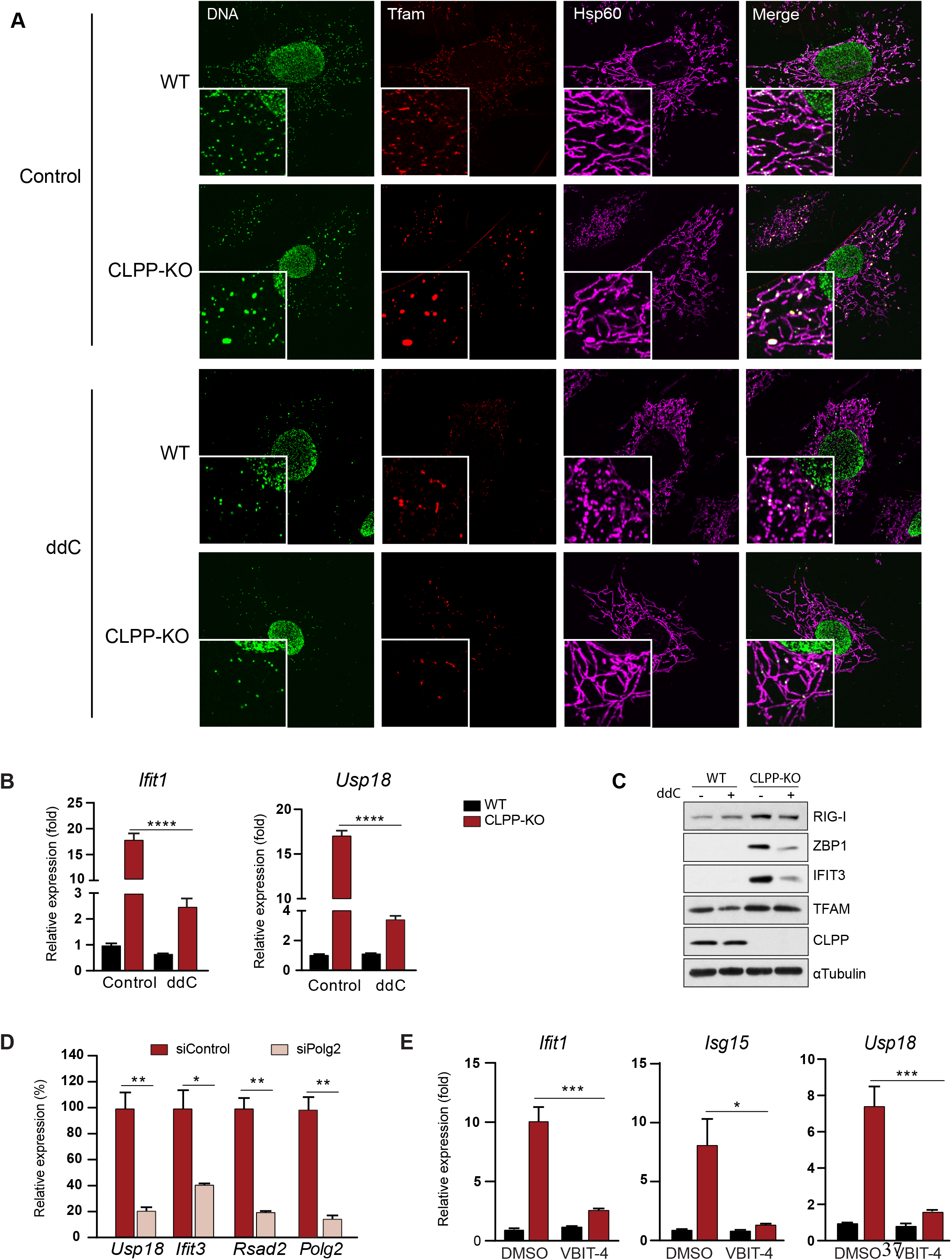
Altered mtDNA homeostasis mediates IFN-I responses in CLPP-KO fibroblasts. A WT and CLPP-KO MEFs were cultured in the presence of ddC (150μM) for 4 days. Cells were harvested after 4 days for confocal microscopy (A) of WT and CLPP-KO MEFs stained with anti-DNA (DNA), anti-Tfam (mtDNA nucleoid marker) and anti-HSP60 (mitochondrial matrix protein). B Quantitative real time PCR of ISG expression after culture in ddC (n=3 MEF lines). C Western blots of ISG protein expression after culture in ddC. D Quantitative real time PCR of ISGs in CLPP-KO MEFs transfected with control or siPolg2 siRNA for 72hrs (n=2 MEF lines). E Quantitative real time PCR of ISGs of WT and CLPP-KO MEFs treated with VBIT-4 (10μM) or Control (DMSO) for 48hrs (n=2 MEF lines). Data information: In D, Real-time PCR data are presented as relative expression percentage (%) considering siControl as 100%. Error bars indicate ± s.e.m. of triplicate technical replicates (Student–8217;s t-test). In B and E, data represented as mean ± s.e.m. of triplicate technical replicates, two-way ANOVA Tukey–8217;s *post-hoc*. *p<0.05, **p<0.01, ****p<0.0001

**Figure 5.**
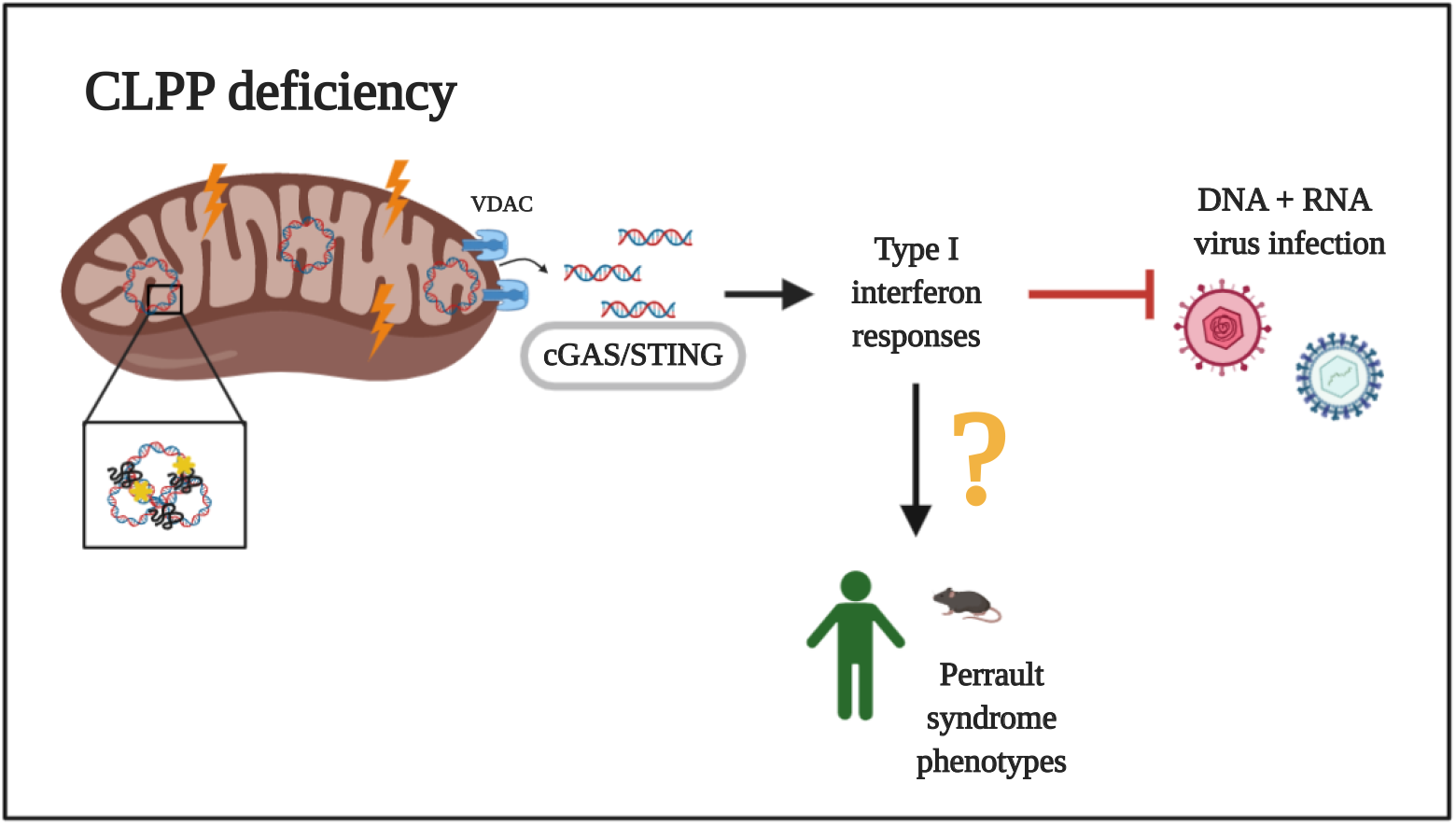
Loss of CLPP activates IFN-I through mtDNA-cGAS-STING-signaling. Absence of CLPP protease leads to mtDNA stress (packaging alterations and cytosolic release), triggering the cGAS-STING-IFN-I pathway, which leads to elevated ISG expression and resistance to DNA and RNA virus infection. Future study should investigate whether chronic activation of this pathway contributes to Perrault syndrome (PS) phenotypes.

Of note, a recent clinical study revealed increased mtDNA copy number and decreased *CLPP* mRNA levels in patient fibroblasts harboring disease-associated *CLPP* mutations (Theunissen et al., 2016). Combined with our results in CLPP deficient cells, these findings collectively suggest a conserved role for CLPP in regulation of mtDNA maintenance and a potential role for mtDNA stress in the pathogenesis of PS. However, our CLPP knockdown and knockout fibroblasts do not exactly phenocopy patient cells harboring *CLPP* missense mutations associated with PS. These disease mutations are noted to alter key amino acids in a cluster near the docking site for CLPX interaction or in the active site of the peptidase (Brodie et al., 2018). Future work to re-introduce synonymous pathogenic mutant CLPP proteins into CLPP-KO mice or to generate germline mutations in *Clpp* via CRISPR/Cas9 will help to determine how these disease relevant mutations contribute to mtDNA instability, IFN-I responses, and disease pathogenesis in PS.

### mtDNA mediates IFN-I responses in CLPP deficient fibroblasts

To next demonstrate that mtDNA is the DAMP triggering ISG responses in CLPP-KO MEFs, we treated cells with dideoxycytidine (ddC), a nucleoside reverse transcriptase inhibitor that specifically inhibits mtDNA replication and causes robust mtDNA depletion with little effect on nuclear DNA (Kasashima et al., 2011). Notably, ddC treatment was sufficient to clear the enlarged/aggregated nucleoids in CLPP-KO MEFs (Fig 4A), as well as robustly reduce mtDNA levels (Fig EV4C). Consistent with the notion that mtDNA stress is driving cGAS-STING-IFN-I signaling in CLPP-KO cells, the pronounced ISG signature observed in CLPP-KO MEFs was markedly reduced at the RNA and protein levels after ddC treatment (Fig 4B and C). In a complementary approach, we transfected MEFs with siRNA against *Polg2* to knockdown the accessory subunit of the mtDNA polymerase pol γ that is required for mtDNA replication (Young and Copeland, 2016). Consistent with ddC results, we observed a significant reduction in ISG transcripts in CLPP-KO cells when *Polg2* was knocked down (Fig 4D).

Recently, it was described that VBIT-4, an inhibitor of voltage-dependent anion channel (VDAC) oligomerization, blocks the formation of mitochondrial pores and prevents the release of mtDNA fragments into the cytosol of *Endog* deficient MEFs (Kim et al., 2019). We next tested this reagent in our MEF lines, and similar to the results with ddC treatment and *Polg2* knockdown, VBIT-4 treatment significantly diminished the strong ISG signature of CLPP-KO MEFs at baseline (Fig 4E). VBIT-4 had no effect on ISG expression induced by recombinant mIFNβ treatment (Fig EV4D), indicating that the abrogation of steady state antiviral responses by VBIT-4 was not due to a general suppression of IFN-I signaling in CLPP-KO cells. Collectively, these results document that mtDNA stress and VDAC-mediated mtDNA release into the cytosol are key triggers of cGAS-STING-IFN-I signaling and antiviral responses in CLPP-KO cells.

While more than 50 nucleoid-associated proteins have been shown to participate in mtDNA maintenance and gene expression, CLPP has not yet been implicated in either process (Lee and Han, 2017). Our findings document that CLPP is necessary for maintaining mtDNA nucleoid organization and distribution, as CLPP null cells exhibit markedly disrupted nucleoid architecture and TFAM aggregation compared to WT cells. While it is unlikely that these effects result from loss of direct CLPP-mtDNA interactions, CLPX was reported to maintain mtDNA nucleoid distribution via TFAM and is involved in mtDNA segregation in human cells (Kasashima et al., 2012). Although CLPX is upregulated in CLPP null cells, we found that downregulation of CLPX by siRNA had no effect on nucleoid aggregation (data not shown) or steady state ISG responses (Fig EV5A). It is also noteworthy that mono-allelic loss of another matrix protease and mtDNA maintenance factor, LONP1, leads to steady state induction of ISGs in MEFs (Key et al., 2020). Collectively, these results suggest that perturbations in mitochondrial matrix protease levels, which disrupt mtDNA homeostasis and stability, generally engage IFN-I signaling, thus linking mammalian UPR^mt^ factors to antiviral innate immunity (Melber and Haynes, 2018; Pellegrino et al., 2014; S. Wang et al., 2018).

Finally, we explored other factors that modulate mtDNA stress and/or signaling to IFN-I in order to rule in or out their involvement in the enhanced ISG signature of CLPP-KO cells. We employed siRNA approaches to target both mitochondrial fusion and fission proteins and examined ISG expression in CLPP-KO MEFs, since mitochondrial dynamics facilitate proper nucleoid distribution and removal of damaged mtDNA (Ban-Ishihara et al., 2013; Malena et al., 2009). However, we found that transient downregulation of mitofusin 1 (Mfn1) or the fission factor dynamin-related protein 1 (Drp1) did not significantly alter the ISG signature in CLPP-KO cells (Fig EV5B). In addition, another study identified ERAL1, a putative 12S rRNA chaperone, as a CLPXP substrate, implicating CLPP in mitoribosomal assembly and mitochondrial translation (Szczepanowska et al., 2016). Although mitoribosomal maturation/assembly was not analyzed carefully in this study, treatment with several concentrations of chloramphenicol (CAM), a mitochondrial translation inhibitor, yielded no effect on the elevated steady state ISG signature of CLPP-KO cells the aberrant nucleoid morphology in CLPP-KO cells (Fig EV5C and data not shown). However, CAM treatment significantly inhibited mitochondrial translation in MEFs, as seen by the significant reductions in cytochrome c oxidase subunit I (mt-COI) protein levels. Future studies aimed at uncovering the molecular mechanisms by which CLPP integrates mitoribosomal assembly, proteostasis, mitochondrial nucleoid organization, and mtDNA release should yield new insight into crosstalk between the mammalian UPR^mt^ and the innate immune system.

In conclusion, Caseinolytic peptidase XP (ClpXP) is an evolutionarily conserved, heteromeric, mitochondrial matrix localized serine protease complex, and despite extensive studies on prokaryotic ClpXP, the diverse functions of CLPXP in mitochondria remain less well defined (Levytskyy et al., 2016). Our results place the cGAS-STING-IFN-I innate immune pathway downstream of CLPP and illuminate new links between mitochondrial proteostasis, mtDNA genome maintenance, and antiviral immunity. Imbalances in mitochondrial protease function, mtDNA damage, and innate immunity are associated with several pathological conditions, such as neurodegenerative disorders, infertility, metabolic syndromes, and cancer (Bárcena et al., 2017; Levytskyy et al., 2016). This work sheds new light on the biology of mammalian CLPP, further advances our understanding of mitochondrial-innate immune crosstalk, and may have implications for understanding PS pathogenesis and other diseases involving CLPP dysregulation.

## Materials and Methods

### Viral stocks and infections

VSV-G/GFP and HSV1/GFP were maintained as described previously (Dalton and Rose, 2001; Desai and Person, 1998). MEFs were plated in 12-well plates at 7×10^4^ cells/well, 16hrs before infection in DMEM+10% FBS. The next morning, virus stocks were diluted at indicated MOIs in serum-free DMEM (D5756-500ML, Sigma). 300 μl serum-free media containing virus (or DMEM alone for the control wells) was gently added to wells. Plates were incubated at 37°C 5% CO_2_ and rocked gently every 15 min for 1hr, after which the supernatant was removed, bleached, and discarded. Then, 0.5 mL of fresh DMEM+10% FBS was added to each well, and the cells were allowed to incubate for the duration of the experiment. At the indicated times post infection, supernatant was collected and cleared by centrifugation at 6,000 rpm, 4°C, then kept at −80°C until further processing. Cells were then fixed and stained for microscopy or lysed in RNA Lysis buffer. For plaque assay: BHK-1 or Vero cells were plated in 6-well plates at 95% confluency in 3% Methylcellulose containing DMEM media. Next, different dilutions of supernatant from VSV or HSV1 infected MEFs were added to the cells and incubated until plaques were visible, between 24-72hrs. Plaques were stained with Cresyl Violet and counted with Image J software.

### Immunofluorescence

Cells were grown on 12mm or 18mm coverslips and treated or infected as described. After washing in PBS, cells were fixed with 4% paraformaldehyde for 20 min, permeabilized with 0.1% Triton X-100 in PBS for 5 min, blocked with PBS containing 10% FBS for 30 min, stained with primary antibodies for 60 min, and stained with secondary antibodies for 60 min. Cells were washed with PBS between each step. Coverslips were mounted with Prolong Gold anti-fade reagent containing DAPI (Molecular Probes). For viral infections, viral GFP fluorescence images were captured using the LionHeart FX fluorescent microscope at 20X and tiling images at 2×2. For mtDNA nucleoid staining, a 60X oil-immersed objective was used and images were taken with a Confocal Laser Scanning Microscope Olympus FV3000. Images were processed in Image J software.

### Quantitative PCR

To measure relative gene expression by qRT-PCR, total cellular RNA was isolated using the Quick-RNA Micro Prep Kit (R1051, Zymo Research). Approximately 250-500ng RNA was normalized across samples and cDNA was generated using the qScript cDNA Synthesis Kit (101414-098, VWR). cDNA was then subjected to qPCR using PerfecTa SYBR Green FastMix (84069, Quantabio) and primers as indicated in Table 1. Three technical replicates were performed for each biological sample, and expression values of each replicate were normalized against Rp137 using the 2 ^−Δddct^ method. For relative expression (fold), control samples were centered at 1; for relative expression (%), control samples were centered at 100%. Mitochondrial DNA abundance analysis was performed as described above using primers specific to mtDNA regions and normalized against nuclear *Tert*.

**Table 1.**
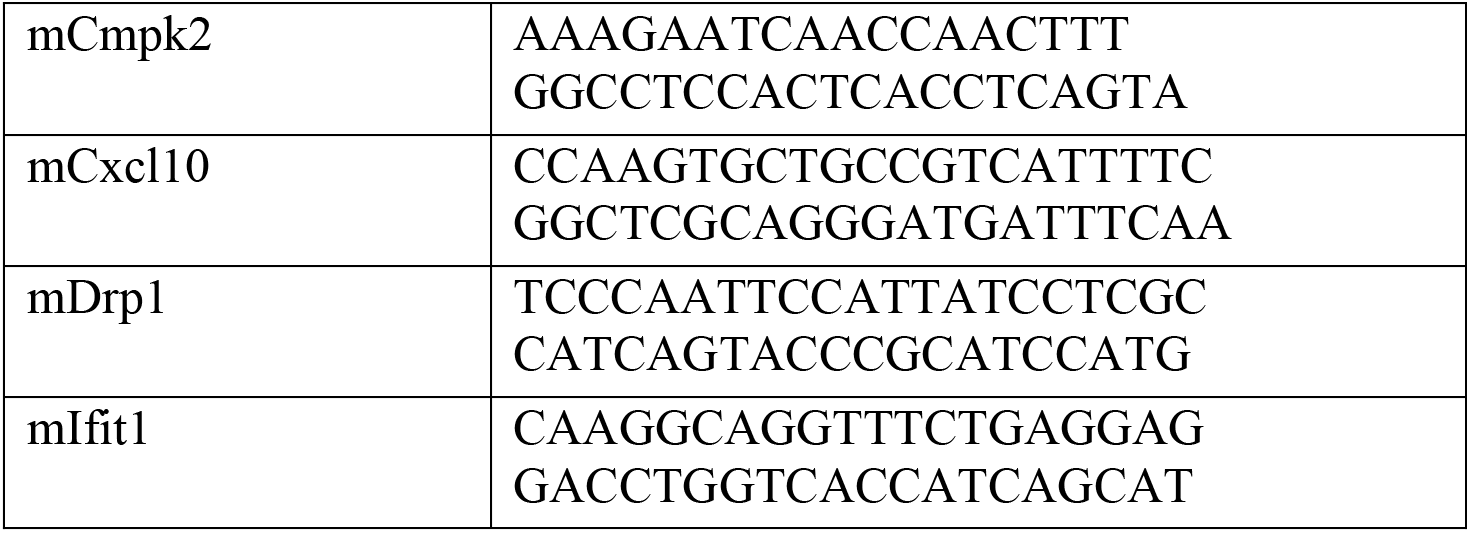
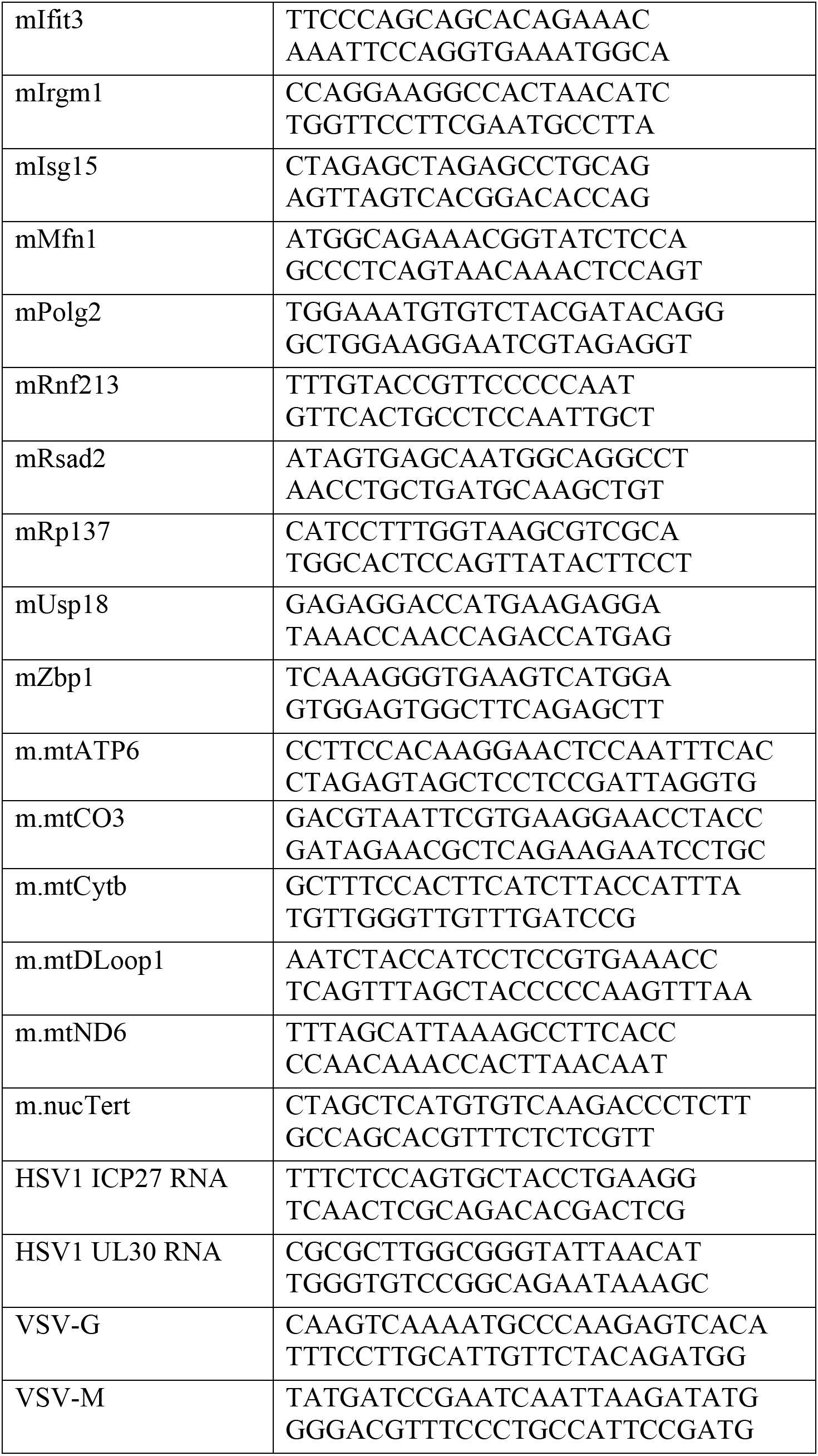
Primer sequences.

### Proteomics

CLPP-KO MEF and their matching controls (n=3) were analyzed by quantitative mass spectrometry essentially as described in Key et al., 2019. Mass spectrometry data were analyzed by MaxQuant (1.5.3.30) (Cox and Mann, 2008) using a mouse proteome Uniprot database (Download 2/2016) and a false discovery rate (FDR) less than 1%. For quantification, proteins were quality filtered according to a minimum of three valid values in one group (n=3) using Perseus software (v. 1.5.2.6). All missing values from this reduced matrix were replaced by background value from normal distribution. For statistical comparison, students t-tests were used. The full datasets will be deposited in the Proteomics Identification Database (PRIDE) upon acceptance of the manuscript.

### Immunoblotting

Proteins from either homogenized tissue (<50mg) or cells (<1×10^7^) were lysed in 1% NP40 buffer, and spun down at 15,000 rpm for 10 min at 4°C. The supernatant was collected as protein lysate and quantified with micro-BCA assay (23235, Proteintech). Between 20-30μg of protein were separated on 10-20% polyacrylamide gradient gels, then transferred onto PVDF membranes at 100V for 1hr. Membranes were dried for 30 min and incubated with primary antibodies overnight at 4°C. Membranes were washed with 1X PBS for 30 min and incubated with HRP-conjugated secondary antibodies for 1hr. Membranes were washed with PBS for 1 hr before developing with Millipore Luminata Crescendo HRP Substrate (WBWR0500).

**Table 2.** List of Antibodies.

| Name | Company | Catalog number | Dilution |
| --- | --- | --- | --- |
| CLPX | Abcam | ab168338 [EP8772] | 1:1,000 |
| IFIT3 | gift from G.Sen at Cleveland Clinic |  | 1:10,000 |
| ZBP1 | Adipogen | AG-20B-0010 |  |
| αTUBULIN | DSHB | 12G10 | 1:5,000 |
| STAT1 | Cell Signaling | 9172S | 1:1,000 |
| cGAS | Cell Signaling | 31659 | 1:1,000 |
| RIG-I | Cell Signaling | 4200S | 1:1,000 |
| IRGM1 | Cell Signaling | 14979 | 1:1,000 |
| MAVS | Cell Signaling | 4983 | 1:1,000 |
| GFP | DSHB | 8H11 | 1:30 |
| TFAM | Millipore | ABE483 | 1:1,000 |
| ANTI-DNA | Millipore | CBL-186 (AC-30-10) | 1:300 |
| IFITM1 | Proteintech | 60074-1-Ig | 1:5,000 |
| IFIT2 | Proteintech | 12604-1 | 1:1,000 |
| CALNEXIN | Proteintech | 10427-2-AP | 1:1,000 |
| β-ACTIN | Proteintech | 66009-1-Ig | 1:5,000 |
| CLPP | Proteintech | 15698-1 | 1:1,000 |
| GAPDH | Proteintech | 60004-1-LG | 1:5,000 |
| HSP60 | Santa Cruz | sc-1052 | 1:5,000 |
| mt-COI | ThermoFisher | 459600 (ID6E1A8) | 1:1,000 |

### Mouse Strains

Embryonal stem cell line IST13563G11 (line G), with heterozygous GeneTrap insertions at *Clpp* intron 2, was re-derived at the Texas Institute for Genomic Medicine (TIGM) on a C57BL/6N background. Heterozygous breeders were mated to generate *Clpp*-null (CLPP-KO) mutant and littermate WT control offspring. *Sting^gt/gt^* (C57BL/6J-*Sting1*^*gt*^/J, strain 017537) and *Ifnar*^*-/-*^ (B6(Cg)-*Ifnar1*^*tm1.2Ees*^/J, strain 028288) were obtained from the Jackson Laboratory. CLPP heterozygous mutant mice were crossed with *Sting*^*gt/gt*^ and *Ifnar*^*-/-*^ for several generations to obtain double KO mice on a C57BL/6NJ mixed background. All animal experiments were conducted in compliance with the guidelines established by the Texas A&M University Institutional Animal Care and Use Committee (IACUC).

### EchoMRI

Body composition analysis on live mice was completed using the EchoMRI™-100H at the Rodent Preclinical Phenotyping Core at Texas A&M University. This analyzer delivers precise body composition measurements of fat, lean, free water, and total water masses in live animals weighing up to 100g.

### Cell Culture

Primary wild-type, CLPP-KO, STING^gt/gt^, IFNAR^-/-^ (IFNAR-KO), CLPP-KO/STING^gt/gt^ and CLPP-KO/IFNAR-KO MEFs were generated from E12.5-14.5 embryos. Cells were maintained in DMEM (D5756-500ML, Sigma) supplemented with 10% FBS (VWR), and sub-cultured for no more than five passages before experiments. Transfection of MEFs with siRNA was performed with Lipo-RNAi Max (13778-150, Invitrogen) in Opti-MEM media (11058021, ThermoFisher Scientific) and the following siRNAs were used at a final concentration of 25nM: sicGAS (MMC.RNAI.N173386.12.1 IDT), siRNA mPolg2 (NM_015810 Sigma-Aldrich), siMfn1 (NM_024200 Sigma-Aldrich), siDrp1 (MMC.RNAI.N152816.12.1), siClpX (MMC.RNAI.N011802.12.2 IDT) and siRNA NTC (Negative Control DsiRNA, 51-01-14-03, IDT) according to manufacturer’s instructions. Cells were harvested for RNA or protein between 48-72hrs later. For Rig-I/MDA5 challenge, cells were transfected with Poly(I:C) (P1530, SIGMA) complexed in polyethylenimine (PEI, 43896, Alfa Aesar) in Opti-MEM media. For LPS, cells were treated with LPS (LPS-B5 Ultrapure, InvivoGen tlrl-pb5lps) at 1μg/ml and harvested for RNA 4hrs later. For mtDNA depletion, ddC (D5782, SIGMA) was resuspended in PBS, added to MEFs at a final concentration of 150uM and refreshed every 48hrs for 4 days. For Chloramphenicol (CAM) treatment, cells were treated with CAM (R4408, SIGMA) at different concentrations (50-200uM) and harvested for protein between 24-48hrs later. For VBIT-4 treatment, VBIT-4 (HY-129122, MedChem Express) was resuspended in DMSO at 10mM and added to the media at a final concentration of 10uM during 48hrs while control cells were treated with DMSO. For VBIT-4 treatment plus challenge, cells were treated with VBIT-4 and 6hrs before the 48hrs harvesting time mIFNb (8234-MB-010/CF R&D) was added to the cells at a final concentration of 1ng/ml. All cells were harvested 48hrs after challenge for RNA analysis. To generate CLPP knockdown human foreskin fibroblasts (HFF-1), MISSION shRNA Lentiviral Transduction Particles against human CLPP (TRCN0000291174) or eGFP (RHS4459) were purchased from Sigma-Aldrich and Horizon Discovery respectively. HFF-1 were transduced with the shRNA encoding lentivirus stocks in the presence of polybrene (8 μg/ml) and stably selected with puromycin.

### Statistical analyses

Error bars displayed throughout the manuscript represent the mean +/− s.e.m. unless otherwise indicated and were calculated from triplicate or quadruplicate technical replicates of each biological sample. Sample sizes were chosen by standard methods to ensure adequate power, and no randomization or blinding was used for animal studies. No statistical method was used to predetermine sample size. Statistical analysis was determined using GraphPad software and statistics tests include: unpaired Student’s t-tests, ordinary one-way ANOVA and two-way ANOVA with Tukey's *post-hoc.* Significance was stablished as *p<0.05; **p<0.01; ***p<0.001; ****p<0.0001; or not significant (ns) when p>0.05. Data shown are representative of 2-3 independent experiments (unless otherwise specified) including microscopy images, Western blots, and viral challenges.

## Acknowledgements

We would like to thank our colleagues in the West Lab for feedback on the manuscript, as well as Dr. Malea Murphy in the Integrated Microscopy and Imaging Laboratory at the Texas A&M College of Medicine for assistance with confocal imaging. S.T.O was partially financed by the Arthur Merx Stiftung Frankfurt. This research was supported by awards W81XWH-17-1-0052 and W81XWH-20-1-0150 to A.P.W. from the Office of the Assistant Secretary of Defense for Health Affairs through the Peer Reviewed Medical Research Programs. Additional support was provided by NIH grant R01HL148153 to A.P.W. and NIEHS P30 ESES029067. Opinions, interpretations, conclusions, and recommendations are those of the author and are not necessarily endorsed by the NIH or Department of Defense.

## Author Contributions

S.T-O. and A.P.W. designed the experiments, analyzed the data, and wrote the manuscript. Y.L., A.M., J.K., and S.M. assisted with experiments, sample processing, and/or data analysis. I.W. performed proteomic analyses. G.A. and S.G. contributed datasets, provided experimental advice, and edited the paper. A.P.W. conceived the project and provided overall direction.

## Expanded View Figure Legends

**Figure EV1.**
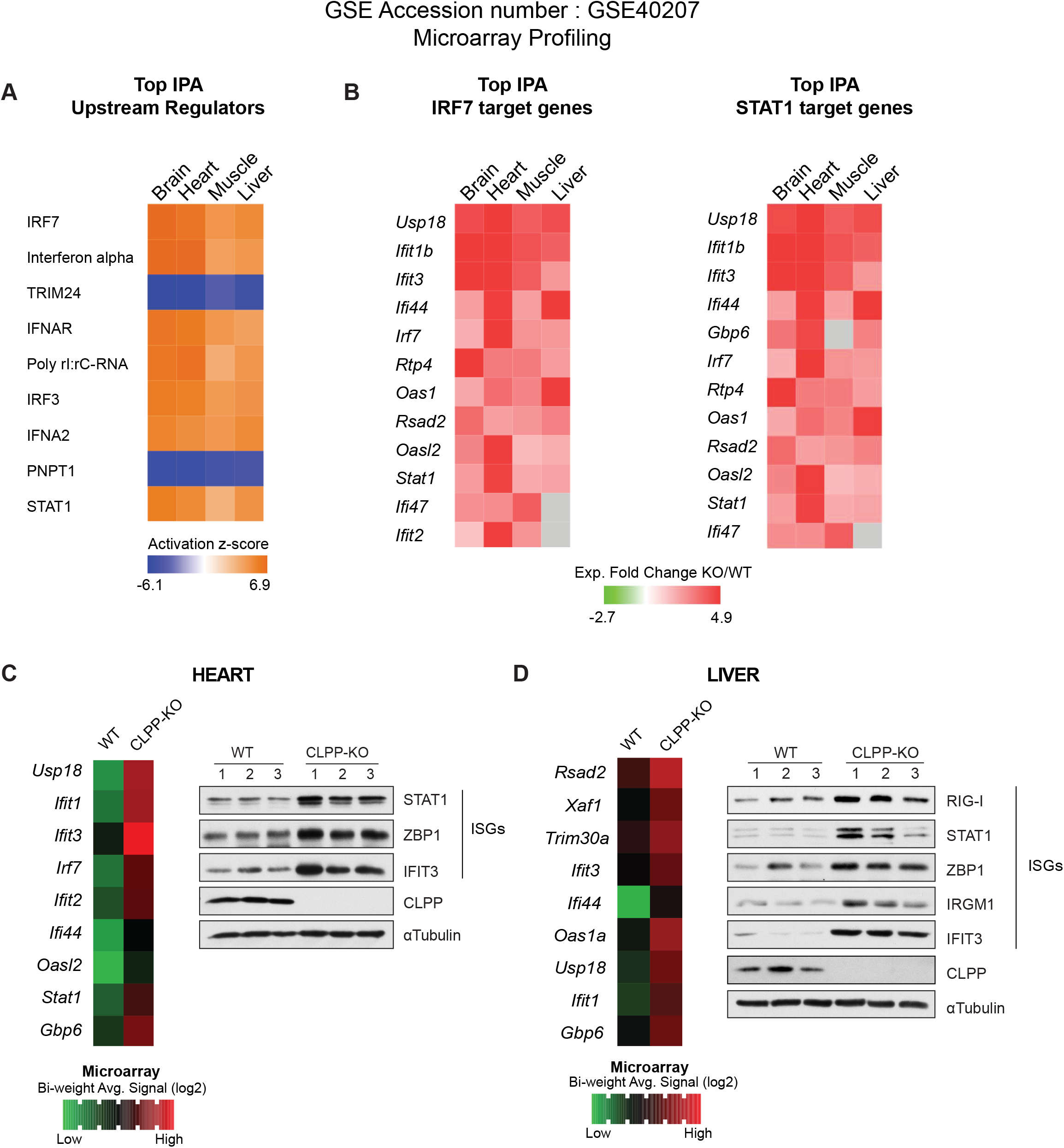
CLPP-KO mice exhibit elevated ISG expression and protein abundance in brain, heart, muscle and liver. A, B Microarray profiling from results reported in Gispert S et al, 2013, GEO Accession GSE40207. Heatmaps of predicted upstream regulators (A) and putative IRF7 and STAT1 target genes of the antiviral gene signature (B) in CLPP-KO brain, heart, muscle and liver tissues using Ingenuity Pathway Analysis (IPA) (n=3). C-D Heatmaps showing most significantly upregulated ISGs from microarray data (GSE40207). Western blots of baseline ISGs in heart (C) and liver (D) tissues of WT and CLPP-KO male mice, between 3-5 months old (n=3).

**Figure EV2.**
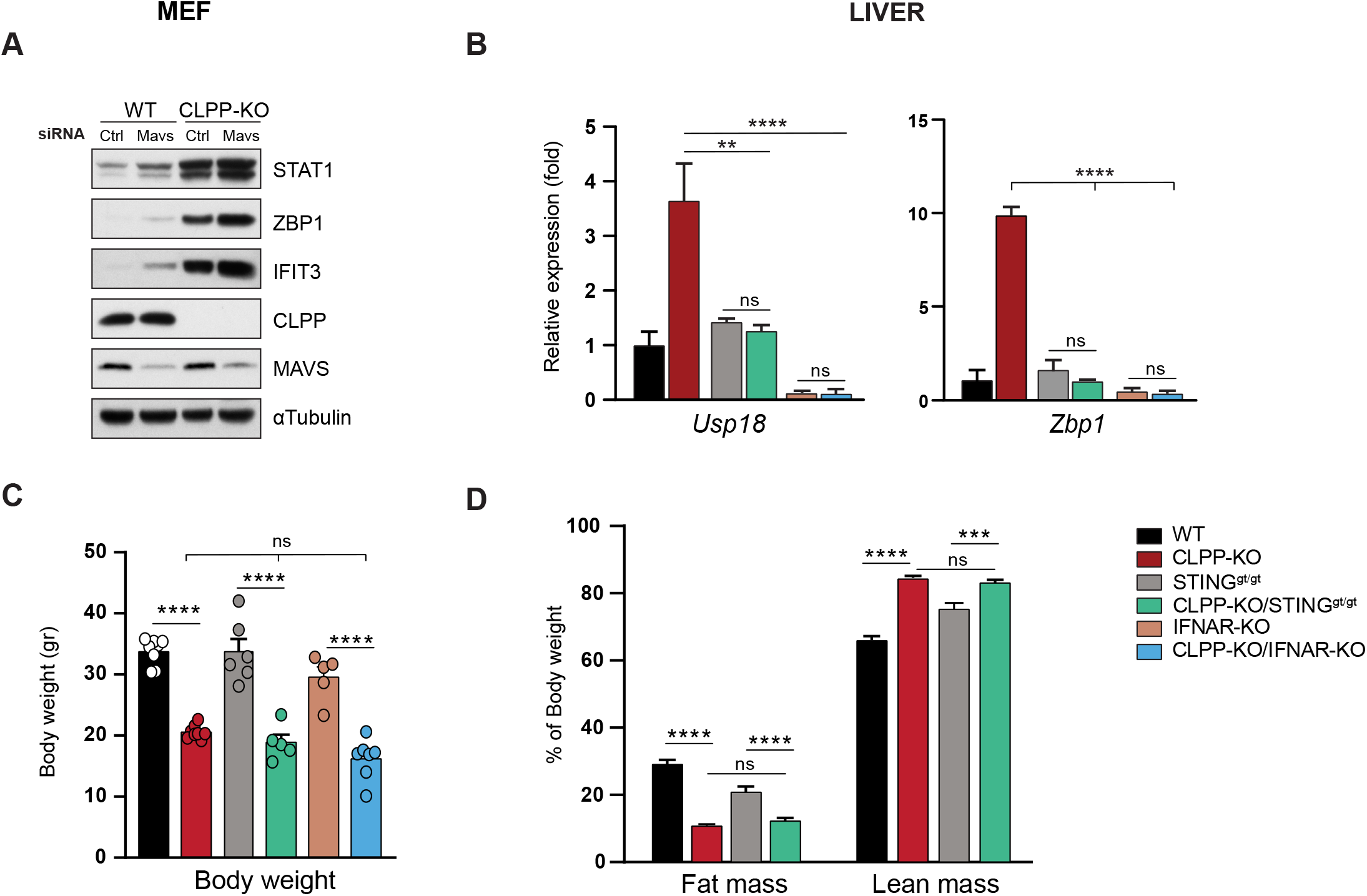
STING and IFNAR signaling drive elevated ISG responses in CLPP deficient cells and tissues, but ablation of IFN-I signaling does not lessen growth retardation of CLPP KO mice. A Western blots of ISGs of WT and CLPP-KO MEFs transfected with siRNA Control (Ctrl) and siMavs for 72hrs. B Quantitative real-time PCR of baseline ISGs of liver tissue from 12-month old male mice (n=3). C Body weight analysis of 5-7-month old male mice from 6 different genotypes (n= 5-8). D Fat mass and lean mass measurements in 7-10-month old female WT, CLPP-KO, STING^gt/gt^ and CLPP-KO/STING^gt/gt^ mice, as assessed by EchoMRI and normalized to body weight (n= 6-10). Data information: In (B-D), data are presented as mean ± s.e.m. of triplicate technical replicates. In (B, C) **p<0.01, ****p<0.0001, ns non-significant (One-way ANOVA Tukey–8217;s *post-hoc*). In D, ***p<0.001, ****p<0.0001, ns non-significant (Two-way ANOVA Tukey–8217;s *post-hoc*).

**Figure EV3.**
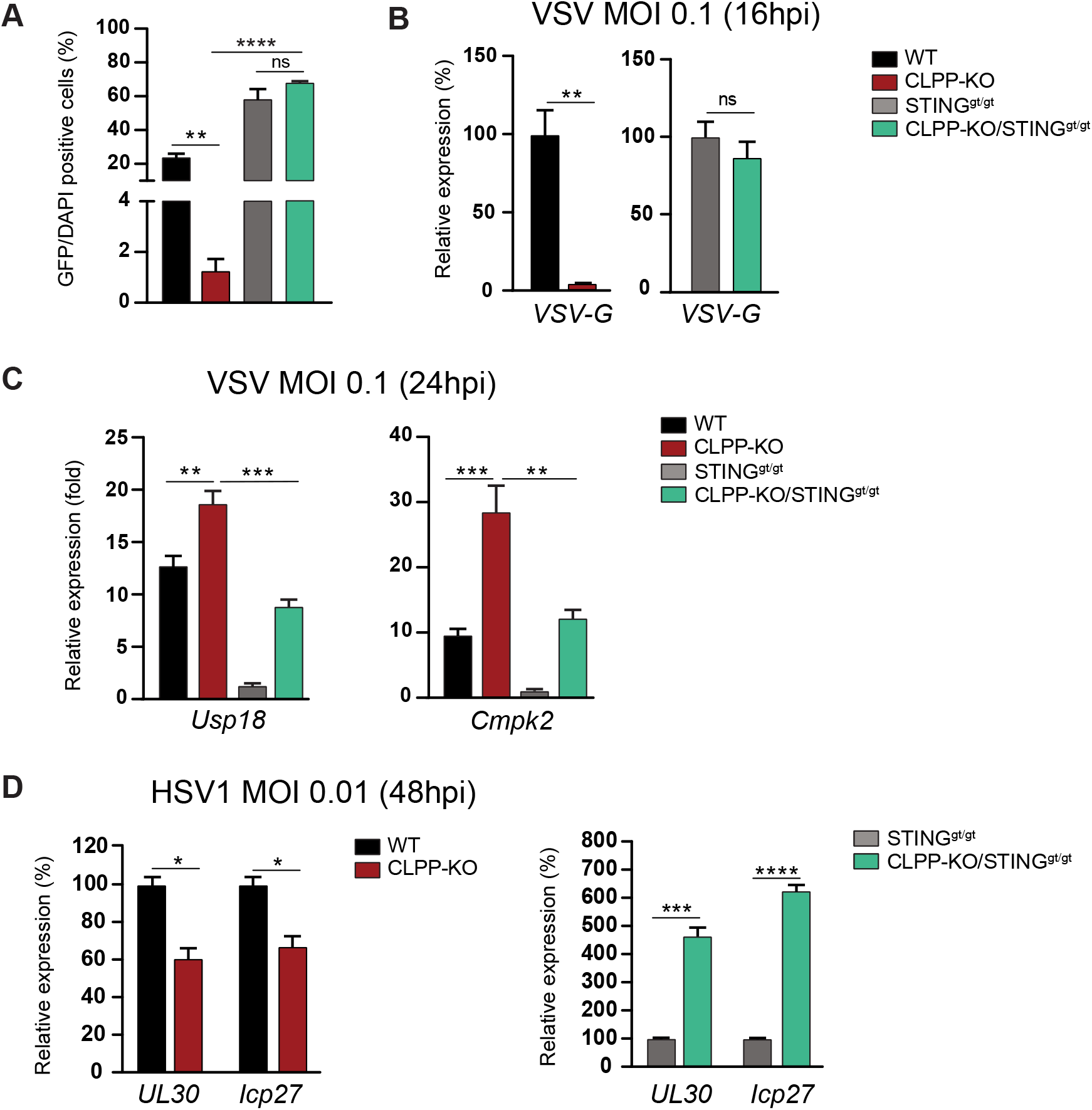
CLPP-KO fibroblasts are more resistant to viral infection owing to elevated activation of the STING pathway. A Quantification of GFP staining in MEFS at MOI 0.1 24hrs after VSV infection. B Quantitative real-time PCR of viral RNA at MOI 0.1 16hrs after VSV infection. C Quantitative real-time PCR of ISGs 24hrs after VSV infection at MOI of 0.1. D Quantitative real-time PCR of viral RNA in MEFs infected with HSV-1 at 48hpi and MOI of 0.01. Data information: In (B, C) Real-time PCR data are presented as relative expression percentage (%) comparing CLPP-KO *vs* WT and CLPP-KO/STING^gt/gt^ *vs* STING^gt/gt^ MEFs. Error bars indicate ± s.e.m. of triplicate technical replicates (Student’s t-test). In (A, C) data are presented as mean ± s.e.m. of triplicate technical replicates (One-way ANOVA Tukey–8217;s *post hoc*). *p<0.05, **p<0.01, ***p<0.001, ****p<0.0001, ns non-significant.

**Figure EV4.**
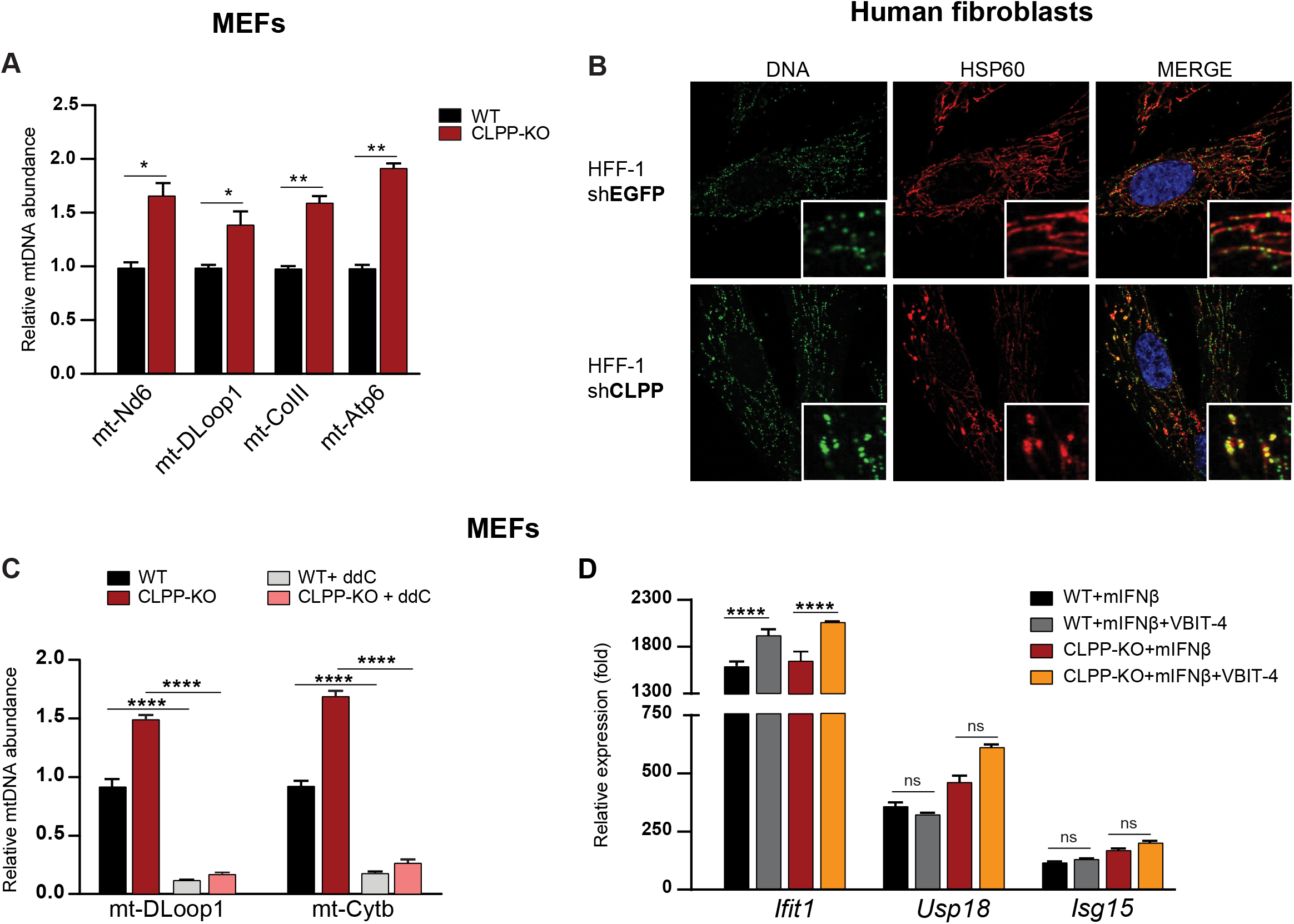
Altered mtDNA homeostasis mediates IFN-I responses in mouse and human CLPP deficient fibroblasts. A Quantification of mtDNA abundance relative to nuclear DNA (Tert) in WT and CLPP-KO MEFs at baseline. B Human foreskin fibroblasts (HFF-1) were transduced with the shRNA encoding CLPP and EGFP (as control) and selected with puromycin (2μg/ml). After selection, cells were plated in 12-well dishes, fixed, and stained with anti-DNA (DNA), anti-HSP60 (mitochondria) and DAPI before imaging on a confocal microscope with a 60X oil immersion objective. C Quantification of mtDNA abundance relative to nuclear DNA (Tert) in WT and CLPP-KO MEF before and after ddC treatment. D Quantitative real time PCR of ISGs of WT and CLPP-KO MEFs treated with VBIT-4 (10μM) and challenged with mIFNβ (1ng/ml) for 6hrs. Data information: In (A, C) data are presented as mean ± s.e.m. of triplicate technical replicates (Student’s test). In D, data are presented as mean ± s.e.m. of triplicate technical replicates (Two-way ANOVA Tukey’s *post hoc*). *p<0.05, **p<0.01, ***p<0.001,****p<0.0001.

**Figure EV5.**
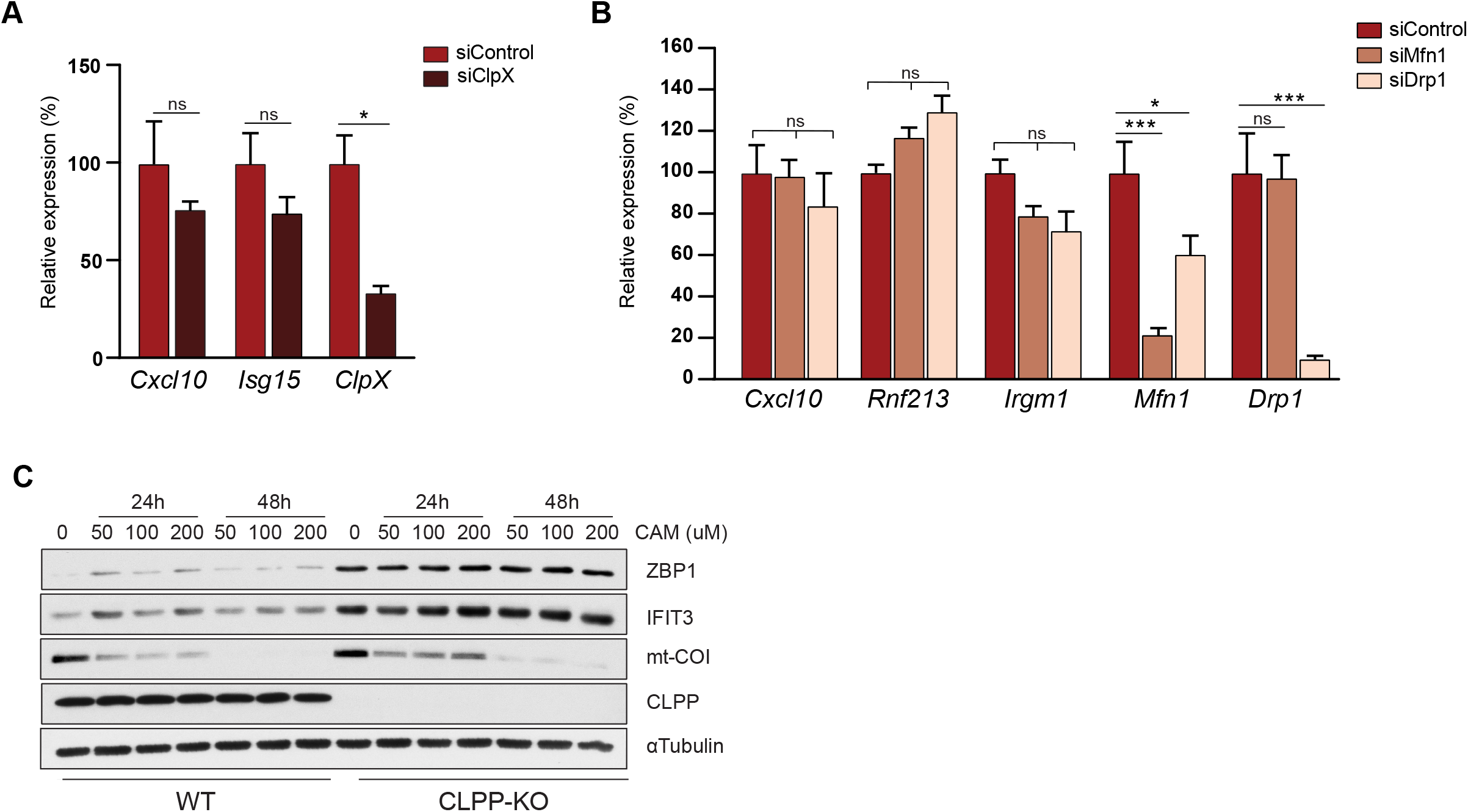
mtDNA stress-mediated IFN-I responses in CLPP-KO fibroblasts are independent of mitochondrial dynamics and translation. A, B Quantitative real time PCR of ISGs in CLPP-KO MEFs transfected with siRNA Control (Ctrl) and (A) siClpX, or (B) siMfn1 and siDrp1 for 72hrs (n=3 MEF lines). C ISG protein levels of WT and CLPP-KO MEFs treated with chloramphenicol (CAM, 50-200μM) over a 24hr to 48hr time course (n=2 MEF lines). Data information: In (A, B) Real-time PCR data are presented as relative expression percentage (%) considering CLPP-KO siControl as 100%. Data are presented as mean ± s.e.m. of triplicate technical replicates. In A, Student’s t-test and in B, One-way ANOVA Tukey’s *post hoc*. *p<0.05, **p<0.01, ***p<0.001, ****p<0.0001, ns non-significant.

## References

Ahn, J., Barber, G.N., 2019. STING signaling and host defense against microbial infection. Exp. Mol. Med. 51, 1–10. https://doi.org/10.1038/s12276-019-0333-0

Asal, S., O. Sobhy, O. Ismail, and E. Bedewy. 2014. Study of the effect of combined interferon and ribavirin therapy on the hearing profile of hepatitis C virus patients. Egypt J Otolaryngol. 31, 4: 237–243.

Baker, T.A., Sauer, R.T., 2012. ClpXP, an ATP-powered unfolding and protein-degradation machine. Biochim. Biophys. Acta 1823, 15–28. https://doi.org/10.1016/j.bbamcr.2011.06.007

Ban-Ishihara, R., Ishihara, T., Sasaki, N., Mihara, K., Ishihara, N., 2013. Dynamics of nucleoid structure regulated by mitochondrial fission contributes to cristae reformation and release of cytochrome c. Proc. Natl. Acad. Sci. U. S. A. 110, 11863–11868. https://doi.org/10.1073/pnas.1301951110

Bárcena, C., P. Mayoral, P. Quiros, and C. López-Otín. 2017. Physiological and Pathological Functions of Mitochondrial Proteases. In: Chakraborti S., Dhalla N. (eds) Proteases in Physiology and Pathology. Springer, Singapore. https://doi.org/10.1007/978-981-10-2513-6_1

Becker, C., Kukat, A., Szczepanowska, K., Hermans, S., Senft, K., Brandscheid, C.P., Maiti, P., Trifunovic, A., 2018. CLPP deficiency protects against metabolic syndrome but hinders adaptive thermogenesis. EMBO Rep. 19: e45126. https://doi.org/10.15252/embr.201745126

Bhandari, V., Wong, K.S., Zhou, J.L., Mabanglo, M.F., Batey, R.A., Houry, W.A., 2018. The Role of ClpP Protease in Bacterial Pathogenesis and Human Diseases. ACS Chem. Biol. 13, 1413–1425. https://doi.org/10.1021/acschembio.8b00124

Bhaskaran, S., Pharaoh, G., Ranjit, R., Murphy, A., Matsuzaki, S., Nair, B.C., Forbes, B., Gispert, S., Auburger, G., Humphries, K.M., Kinter, M., Griffin, T.M., Deepa, S.S., 2018. Loss of mitochondrial protease ClpP protects mice from diet-induced obesity and insulin resistance. EMBO Rep. 19: e45009. https://doi.org/10.15252/embr.201745009

Brodie, E.J., Zhan, H., Saiyed, T., Truscott, K.N., Dougan, D.A., 2018. Perrault syndrome type 3 caused by diverse molecular defects in CLPP. Sci. Rep. 8, 12862. https://doi.org/10.1038/s41598-018-30311-1

Corcoran, J.A., Saffran, H.A., Duguay, B.A., Smiley, J.R., 2009. Herpes simplex virus UL12.5 targets mitochondria through a mitochondrial localization sequence proximal to the N terminus. J. Virol. 83, 2601–2610. https://doi.org/10.1128/JVI.02087-08

Cox, J., Mann, M., 2008. MaxQuant enables high peptide identification rates, individualized p.p.b.-range mass accuracies and proteome-wide protein quantification. Nat. Biotechnol. 26, 1367–1372. https://doi.org/10.1038/nbt.1511

Dalton, K.P., Rose, J.K., 2001. Vesicular stomatitis virus glycoprotein containing the entire green fluorescent protein on its cytoplasmic domain is incorporated efficiently into virus particles. Virology 279, 414–421. https://doi.org/10.1006/viro.2000.0736

Deepa, S.S., Bhaskaran, S., Ranjit, R., Qaisar, R., Nair, B.C., Liu, Y., Walsh, M.E., Fok, W.C., Van Remmen, H., 2016. Down-regulation of the mitochondrial matrix peptidase ClpP in muscle cells causes mitochondrial dysfunction and decreases cell proliferation. Free Radic. Biol. Med. 91, 281–292. https://doi.org/10.1016/j.freeradbiomed.2015.12.021

Desai, P., Person, S., 1998. Incorporation of the green fluorescent protein into the herpes simplex virus type 1 capsid. J. Virol. 72, 7563–7568.

Dhir, A., Dhir, S., Borowski, L.S., Jimenez, L., Teitell, M., Rötig, A., Crow, Y.J., Rice, G.I., Duffy, D., Tamby, C., Nojima, T., Munnich, A., Schiff, M., de Almeida, C.R., Rehwinkel, J., Dziembowski, A., Szczesny, R.J., Proudfoot, N.J., 2018. Mitochondrial double-stranded RNA triggers antiviral signalling in humans. Nature 560, 238–242. https://doi.org/10.1038/s41586-018-0363-0

Duarte, L.F., Farías, M.A., Álvarez, D.M., Bueno, S.M., Riedel, C.A., González, P.A., 2019. Herpes Simplex Virus Type 1 Infection of the Central Nervous System: Insights Into Proposed Interrelationships With Neurodegenerative Disorders. Front. Cell. Neurosci. 13, 36. https://doi.org/10.3389/fncel.2019.00046

Formann, E., Stauber, R., Denk, D.-M., Jessner, W., Zollner, G., Munda-Steindl, P., Gangl, A., Ferenci, P., 2004. Sudden Hearing Loss in Patients with Chronic Hepatitis C Treated with Pegylated Interferon/Ribavirin. Am. J. Gastroenterol. 99, 873–877. https://doi.org/10.1111/j.1572-0241.2004.30372.x

Franz, K.M., Neidermyer, W.J., Tan, Y.-J., Whelan, S.P.J., Kagan, J.C., 2018. STING-dependent translation inhibition restricts RNA virus replication. Proc. Natl. Acad. Sci. 115, E2058–E2067. https://doi.org/10.1073/pnas.1716937115

Gispert, S., Parganlija, D., Klinkenberg, M., Dröse, S., Wittig, I., Mittelbronn, M., Grzmil, P., Koob, S., Hamann, A., Walter, M., Büchel, F., Adler, T., Hrabé de Angelis, M., Busch, D.H., Zell, A., Reichert, A.S., Brandt, U., Osiewacz, H.D., Jendrach, M., Auburger, G., 2013. Loss of mitochondrial peptidase Clpp leads to infertility, hearing loss plus growth retardation via accumulation of CLPX, mtDNA and inflammatory factors. Hum. Mol. Genet. 22, 4871–4887. https://doi.org/10.1093/hmg/ddt338

Graves, P.R., Aponte-Collazo, L.J., Fennell, E.M.J., Graves, A.C., Hale, A.E., Dicheva, N., Herring, L.E., Gilbert, T.S.K., East, M.P., McDonald, I.M., Lockett, M.R., Ashamalla, H., Moorman, N.J., Karanewsky, D.S., Iwanowicz, E.J., Holmuhamedov, E., Graves, L.M., 2019. Mitochondrial Protease ClpP is a Target for the Anticancer Compounds ONC201 and Related Analogues. ACS Chem. Biol. 14, 1020–1029. https://doi.org/10.1021/acschembio.9b00222

Haynes, C.M., Petrova, K., Benedetti, C., Yang, Y., Ron, D., 2007. ClpP mediates activation of a mitochondrial unfolded protein response in C. elegans. Dev. Cell 13, 467–480. https://doi.org/10.1016/j.devcel.2007.07.016

Hekman, A.C., Trapman, J., Mulder, A.H., van Gaalen, J.L., Zwarthoff, E.C., 1988. Interferon expression in the testes of transgenic mice leads to sterility. J. Biol. Chem. 263, 12151–12155.

Ishizawa, J., Zarabi, S.F., Davis, R.E., Halgas, O., Nii, T., Jitkova, Y., Zhao, R., St-Germain, J., Heese, L.E., Egan, G., Ruvolo, V.R., Barghout, S.H., Nishida, Y., Hurren, R., Ma, W., Gronda, M., Link, T., Wong, K., Mabanglo, M., Kojima, K., Borthakur, G., MacLean, N., Ma, M.C.J., Leber, A.B., Minden, M.D., Houry, W., Kantarjian, H., Stogniew, M., Raught, B., Pai, E.F., Schimmer, A.D., Andreeff, M., 2019. Mitochondrial ClpP-Mediated Proteolysis Induces Selective Cancer Cell Lethality. Cancer Cell 35, 721–737.e9. https://doi.org/10.1016/j.ccell.2019.03.014

Iwakura, Y., Asano, M., Nishimune, Y., Kawade, Y., 1988. Male sterility of transgenic mice carrying exogenous mouse interferon-beta gene under the control of the metallothionein enhancer-promoter. EMBO J. 7, 3757–3762. https://doi.org/10.1002/j.1460-2075.1988.tb03259.x

Jenkinson, E.M., Rehman, A.U., Walsh, T., Clayton-Smith, J., Lee, K., Morell, R.J., Drummond, M.C., Khan, S.N., Naeem, M.A., Rauf, B., Billington, N., Schultz, J.M., Urquhart, J.E., Lee, M.K., Berry, A., Hanley, N.A., Mehta, S., Cilliers, D., Clayton, P.E., Kingston, H., Smith, M.J., Warner, T.T., University of Washington Center for Mendelian Genomics, Black, G.C., Trump, D., Davis, J.R.E., Ahmad, W., Leal, S.M., Riazuddin, S., King, M.-C., Friedman, T.B., Newman, W.G., 2013. Perrault syndrome is caused by recessive mutations in CLPP, encoding a mitochondrial ATP-dependent chambered protease. Am. J. Hum. Genet. 92, 605–613. https://doi.org/10.1016/j.ajhg.2013.02.013

Kasashima, K., Sumitani, M., Endo, H., 2012. Maintenance of mitochondrial genome distribution by mitochondrial AAA+ protein ClpX. Exp. Cell Res. 318, 2335–2343. https://doi.org/10.1016/j.yexcr.2012.07.012

Kasashima, K., Sumitani, M., Endo, H., 2011. Human mitochondrial transcription factor A is required for the segregation of mitochondrial DNA in cultured cells. Exp. Cell Res. 317, 210–220. https://doi.org/10.1016/j.yexcr.2010.10.008

Key, J., Kohli, A., Bárcena, C., López-Otín, C., Heidler, J., Wittig, I., Auburger, G., 2019. Global Proteome of LonP1+/− Mouse Embryonal Fibroblasts Reveals Impact on Respiratory Chain, but No Interdependence between Eral1 and Mitoribosomes. Int. J. Mol. Sci. 20, 4523. https://doi.org/10.3390/ijms20184523

Key, J., Maletzko, A., Kohli, A., Gispert, S., Torres-Odio, S., Wittig, I., Heidler, J., Bárcena, C., López-Otín, C., Lei, Y., West, A.P., Münch, C., Auburger, G., 2020. Loss of mitochondrial ClpP, Lonp1, and Tfam triggers transcriptional induction of Rnf213, a susceptibility factor for moyamoya disease. Neurogenetics 21, 187–203. https://doi.org/10.1007/s10048-020-00609-2

Kim, J., Gupta, R., Blanco, L.P., Yang, S., Shteinfer-Kuzmine, A., Wang, K., Zhu, J., Yoon, H.E., Wang, X., Kerkhofs, M., Kang, H., Brown, A.L., Park, S.-J., Xu, X., van Rilland, E.Z., Kim, M.K., Cohen, J.I., Kaplan, M.J., Shoshan-Barmatz, V., Chung, J.H., 2019. VDAC oligomers form mitochondrial pores to release mtDNA fragments and promote lupus-like disease. Science 366, 1531–1536. https://doi.org/10.1126/science.aav4011

Lee, S.R., Han, J., 2017. Mitochondrial Nucleoid: Shield and Switch of the Mitochondrial Genome. Oxid. Med. Cell. Longev. https://doi.org/10.1155/2017/8060949

Levytskyy, R.M., Germany, E.M., Khalimonchuk, O., 2016. Mitochondrial Quality Control Proteases in Neuronal Welfare. J. Neuroimmune Pharmacol. Off. J. Soc. NeuroImmune Pharmacol. 11, 629–644. https://doi.org/10.1007/s11481-016-9683-8

Malena, A., Loro, E., Di Re, M., Holt, I.J., Vergani, L., 2009. Inhibition of mitochondrial fission favours mutant over wild-type mitochondrial DNA. Hum. Mol. Genet. 18, 3407–3416. https://doi.org/10.1093/hmg/ddp281

Melber, A., Haynes, C.M., 2018. UPRmt regulation and output: a stress response mediated by mitochondrial-nuclear communication. Cell Res. 28, 281–295. https://doi.org/10.1038/cr.2018.16

Nakahira, K., Hisata, S., Choi, A.M.K., 2015. The Roles of Mitochondrial Damage-Associated Molecular Patterns in Diseases. Antioxid. Redox Signal. 23, 1329–1350. https://doi.org/10.1089/ars.2015.6407

Pellegrino, M.W., Nargund, A.M., Kirienko, N.V., Gillis, R., Fiorese, C.J., Haynes, C.M., 2014. Mitochondrial UPR-regulated innate immunity provides resistance to pathogen infection. Nature 516, 414–417. https://doi.org/10.1038/nature13818

Quirós, P.M., Langer, T., López-Otín, C., 2015. New roles for mitochondrial proteases in health, ageing and disease. Nat. Rev. Mol. Cell Biol. 16, 345–359. https://doi.org/10.1038/nrm3984

Reshi, L., Wang, H.-V., Hong, J.-R., 2018. Modulation of Mitochondria During Viral Infections. Mitochondrial Dis. E. Taskin, C. Guven and Y. Sevgiler, IntechOpen. https://doi.org/10.5772/intechopen.73036

Rongvaux, A., Jackson, R., Harman, C.C.D., Li, T., West, A.P., de Zoete, M.R., Wu, Y., Yordy, B., Lakhani, S.A., Kuan, C.-Y., Taniguchi, T., Shadel, G.S., Chen, Z.J., Iwasaki, A., Flavell, R.A., 2014. Apoptotic caspases prevent the induction of type I interferons by mitochondrial DNA. Cell 159, 1563–1577. https://doi.org/10.1016/j.cell.2014.11.037

Saffran, H.A., Pare, J.M., Corcoran, J.A., Weller, S.K., Smiley, J.R., 2007. Herpes simplex virus eliminates host mitochondrial DNA. EMBO Rep. 8, 188–193. https://doi.org/10.1038/sj.embor.7400878

Szczepanowska, K., Maiti, P., Kukat, A., Hofsetz, E., Nolte, H., Senft, K., Becker, C., Ruzzenente, B., Hornig-Do, H.-T., Wibom, R., Wiesner, R.J., Krüger, M., Trifunovic, A., 2016. CLPP coordinates mitoribosomal assembly through the regulation of ERAL1 levels. EMBO J. 35, 2566–2583. https://doi.org/10.15252/embj.201694253

Theunissen, T.E.J., Szklarczyk, R., Gerards, M., Hellebrekers, D.M.E.I., Mulder-Den Hartog, E.N.M., Vanoevelen, J., Kamps, R., de Koning, B., Rutledge, S.L., Schmitt-Mechelke, T., van Berkel, C.G.M., van der Knaap, M.S., de Coo, I.F.M., Smeets, H.J.M., 2016. Specific MRI Abnormalities Reveal Severe Perrault Syndrome due to CLPP Defects. Front. Neurol. 7, 203. https://doi.org/10.3389/fneur.2016.00203

Ulusoy, E., Çayan, S., Yilmaz, N., Aktaş, S., Acar, D., Doruk, E., 2004. Interferon α-2b may impair testicular histology including spermatogenesis in a rat model. Arch. Androl. 50, 379–385. https://doi.org/10.1080/01485010490474823

Wang, S., Gao, K., Liu, Y., 2018. UPRmt coordinates immunity to maintain mitochondrial homeostasis and animal fitness. Mitochondrion, Mitochondria in Innate and Adaptive Immunity 41, 9–13. https://doi.org/10.1016/j.mito.2017.11.004

Wang, T., Babayev, E., Jiang, Z., Li, G., Zhang, M., Esencan, E., Horvath, T., Seli, E., 2018. Mitochondrial unfolded protein response gene Clpp is required to maintain ovarian follicular reserve during aging, for oocyte competence, and development of pre-implantation embryos. Aging Cell 17, e12784. https://doi.org/10.1111/acel.12784

West, A.P., 2017. Mitochondrial dysfunction as a trigger of innate immune responses and inflammation. Toxicology 391, 54–63. https://doi.org/10.1016/j.tox.2017.07.016

West, A.P., Khoury-Hanold, W., Staron, M., Tal, M.C., Pineda, C.M., Lang, S.M., Bestwick, M., Duguay, B.A., Raimundo, N., MacDuff, D.A., Kaech, S.M., Smiley, J.R., Means, R.E., Iwasaki, A., Shadel, G.S., 2015. Mitochondrial DNA stress primes the antiviral innate immune response. Nature 520, 553–557. https://doi.org/10.1038/nature14156

West, A.P., Shadel, G.S., 2017. Mitochondrial DNA in innate immune responses and inflammatory pathology. Nat. Rev. Immunol. 17, 363–375. https://doi.org/10.1038/nri.2017.21

Youle, R.J., 2019. Mitochondria–8212;Striking a balance between host and endosymbiont. Science 365. https://doi.org/10.1126/science.aaw9855

Young, M.J., Copeland, W.C., 2016. Human mitochondrial DNA replication machinery and disease. Curr. Opin. Genet. Dev. 38, 52–62. https://doi.org/10.1016/j.gde.2016.03.005

